# Vestibular CCK signaling drives motion-induced malaise

**DOI:** 10.1101/2021.09.08.459510

**Authors:** Pablo Machuca-Márquez, Laura Sánchez-Benito, Fabien Menardy, Andrea Urpi, Mònica Girona, Emma Puighermanal, Isabella Appiah, Richard Palmiter, Elisenda Sanz, Albert Quintana

## Abstract

Travel can induce motion sickness (MS) in susceptible individuals. MS is an evolutionary conserved mechanism caused by mismatches between motion-related sensory information and past visual and motion memory, triggering a malaise accompanied by hypolocomotion, hypothermia, hypophagia and nausea. Vestibular nuclei (VN) are critical for the processing of movement input from the inner ear. Motion-induced activation of VN neurons recapitulates MS-related signs. However, the genetic identity of VN neurons mediating MS- related autonomic and aversive responses remains unknown. Here, we identify a central role of cholecystokinin (CCK)-expressing VN neurons in motion-induced malaise. Moreover, we show that CCK VN inputs onto the parabrachial nucleus activate *Calca*- expressing neurons and are sufficient to establish avoidance to novel food, which is prevented by CCK-A receptor antagonism. These observations provide greater insight into the neurobiological regulation of MS by identifying the neural substrates of MS and providing potential targets for treatment.

**SIGNIFICANCE STATEMENT:** We live in an age where travel is paramount. However, one of the most disabling conditions inherent to traveling is motion sickness (MS). While studies have underscored the role of the vestibular system in the development of MS, the neuronal populations involved in motion-induced malaise remain largely unknown.

Here, we describe the vestibular pathways eliciting MS responses, and identify a key role for cholecystokinin (CCK)-expressing vestibular neurons. We reveal that a vestibulo-parabrachial (PBN) CCKergic projection is sufficient to induce conditioned taste aversion, likely through the activation of calcitonin gene-related peptide-expressing PBN neurons. Finally, we underscore the role of CCK-A receptor signaling as a novel druggable target to treat MS, providing novel insight on the neurobiological substrates of MS.

## INTRODUCTION

Motion sickness (MS) is an unpleasant autonomic physiological condition that occurs in healthy individuals undergoing passive or even illusory motion. MS signs and symptoms include pallor, cold sweating, yawning, retching and vomiting, vertigo, anorexia, drowsiness, and even severe pain (1–5). MS is highly conserved among species (4, 6–9). Thus, while still debated (10), it has been posited that MS may be the byproduct of an evolutionary mechanism acting as an early toxin-ingestion warning system leading to reduced metabolism (hypothermia and drowsiness), expulsion of the toxin (vomiting) and future avoidance of the ingested substance (6).

It is widely accepted that MS arises from conflicts between actual visual/vestibular sensory inputs and the expected motion and body position information based on past memories (11–13). Accordingly, the necessity of a functional vestibular system for the development of MS was identified long ago (14, 15). Movement-related information processed by the vestibular organ in the inner ear is relayed directly to the medullo-pontine vestibular nuclei (VN). Compelling evidence demonstrates that VN neurons are central to MS neurobiological regulation by showing that activation of VN neurons through provocative motion reproduced MS-like autonomic alterations in rats and mice (16–18). In addition, diseases that affect VN function are associated to autonomic dysregulation such as vertigo, nausea, and vomiting (19).

The role of the VN in the control of the body orientation system, mainly by means of vestibular reflexes at the ocular, head, neck, and spinal levels, has been well described (20). Furthermore, vestibular outputs have been shown to modulate blood pressure with posture change (21, 22). However, how vestibular function governs motion-induced changes in other MS-mediated responses, such as hypolocomotion, appetite suppression, loss of body temperature, or acquisition of a conditioned taste avoidance (CTA) are still unknown (3, 4, 8, 16, 17, 23–25). Mapping of neuronal activity after provocative motion has identified potential neural substrates for vestibular-induced physiological responses, such as the nucleus of the solitary tract (NTS), paraventricular nucleus of the hypothalamus (PVN), parabrachial nucleus (PBN), central amygdala, dorsal raphe nucleus, locus coeruleus and area postrema (16, 17). Among them, both the NTS and PBN are known to process visceral sensory input in the brainstem (26) and receive direct connections from the VN (27–29).

Excitatory neurons are the main projection-neuron type in the VN (30) and have been suggested to participate in autonomic responses after hypergravity (31) and postural imbalance (32). Furthermore, provocative-motion stimuli activate glutamatergic vesicular glutamate transporter 2 (VGLUT2)-expressing VN neurons that in turn project axons to nuclei such as the PBN (33–35). Thus, we hypothesized that genetically defined, glutamatergic neuronal (sub)populations and circuits in the VN would be sufficient and/or necessary to develop MS-induced autonomic regulation and/or aversive learning. To that end, we used cell-type-specific transcriptomics combined with optogenetic and chemogenetic approaches to identify the underlying vestibular circuitry of MS-induced responses.

## RESULTS

### Glutamatergic vestibular neurons sustain motion-induced autonomic responses

Assessment of MS in mice is hindered by the lack of emetic reflex and difficulty to unequivocally identify nausea (5, 8, 36). However, behavioral and autonomic signs of MS are consistently observed in mice subjected to rotatory or gravitational paradigms (8, 16–18). Thus, we sought to establish a rotational paradigm sufficient to develop robust MS-like symptoms (Figure 1A). Four repeated 1-min, 4-*g* accelerations (spin stimulation) consistently induced transient hypolocomotion (Figure 1B and S1A). Furthermore, a significant decrease in food intake was observed in food-deprived mice after spin stimulation compared to controls (Figure 1C), in line with the appetite suppression observed in MS (3, 4, 8, 16, 18, 24). In addition, a decrease in core body temperature, another classical MS response (3, 4, 8, 17, 24, 25), was observed after the rotational stimulus compared to controls, with a maximum temperature difference of 4°C that occurred ∼17.5 min after rotation onset (Figure 1D).

**Figure 1.**
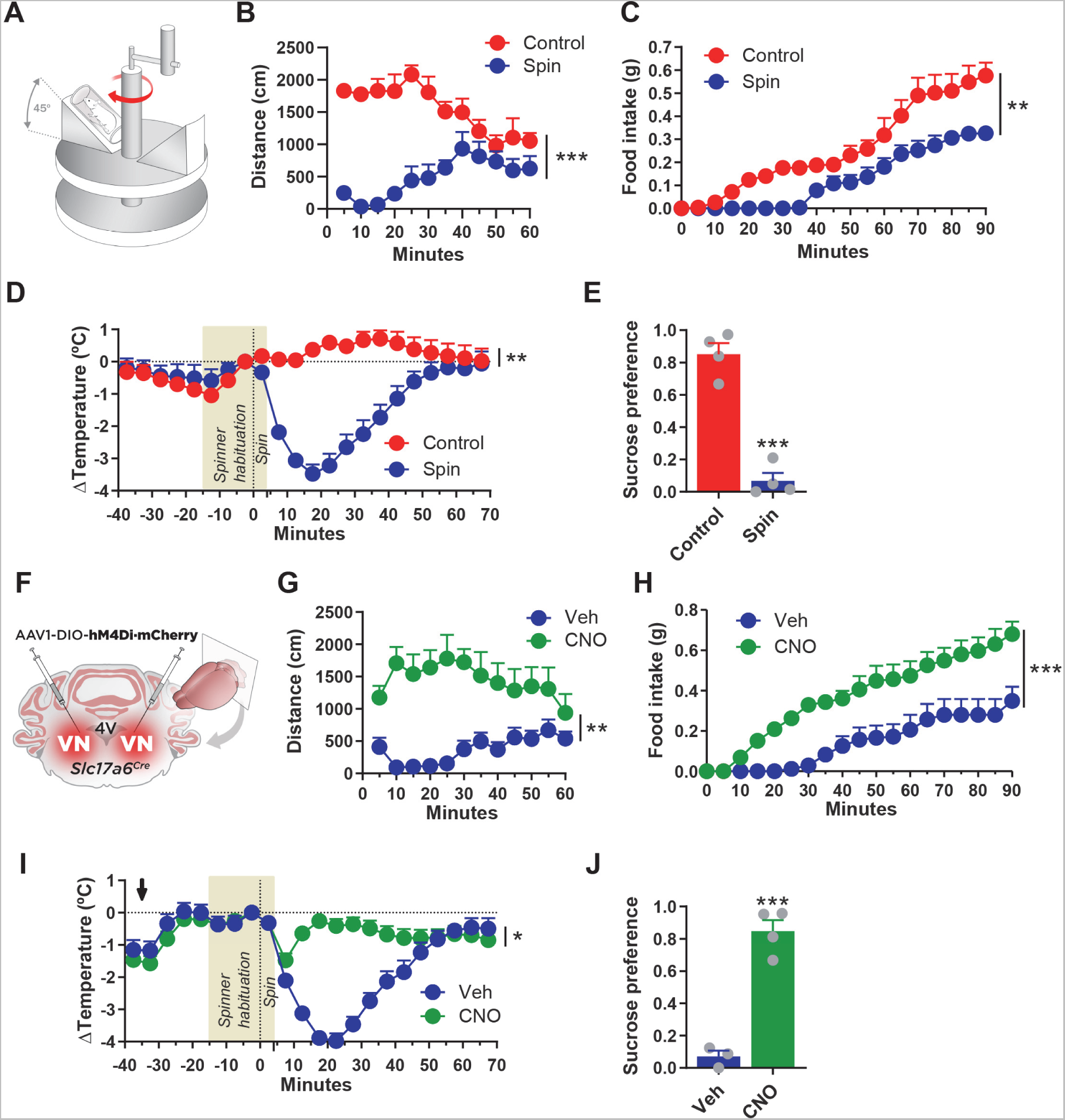
VGLUT2^VN^ neuron inactivation blocks rotation-induced kinetosis in mice. (A) Mice were subjected to a rotational stimulus using a custom-made rotary device. Perpendicular distance from the mouse head to the axis of rotation was 5 cm. Mouse body inclination angle was 45°. (B) Traveled distance during a 60-min open-field test after spin or control stimulation (n=6; two-way ANOVA, *P* <0.001 effect of spin). (C) Cumulative food intake after spin or control stimulation (n=6; two-way ANOVA, *P*<0.01 effect of spin). Mice were food deprived for 24 h prior to the test. (D) Body temperature difference (ΔT) after spin or control stimulation (n=5; two-way ANOVA, *P*<0.01 effect of spin). (E) Conditioned taste avoidance (CTA) response in mice exposed to a two-bottle-based test pairing a 5% sucrose solution to rotational stimulus (n=4; t-test, *P*<0.001). (F) *Slc17a6^Cre^* mice were bilaterally injected in the vestibular nuclei (VN) with AAV1-DIO-hM4DiꞏmCherry (VGLUT2^VN^:hM4Di mice) to inhibit glutamatergic neurons upon CNO administration. (G) Total distance traveled during a 60-min open-field session after spin stimulation in CNO- or vehicle-injected VGLUT2^VN^:hM4Di mice (n=10; two-way ANOVA, *P*<0.01 effect of drug). (H) Cumulative food intake after spin stimulation in CNO- or vehicle-injected VGLUT2^VN^:hM4Di mice (n=7; two-way ANOVA, *P*<0.001 effect of drug). Mice were food deprived for 24 h prior to the test. (I) Core body temperature difference (ΔT) after spin stimulation in CNO- or vehicle-injected VGLUT2^VN^:hM4Di mice (n=4; two-way ANOVA, *P*<0.05 effect of drug). The arrow shows the time of injection. (J) CTA response in mice exposed to a two-bottle-based test pairing a 5% sucrose solution to rotational stimulus in VGLUT2^VN^:hM4Di mice injected with CNO (n=4) or vehicle (n=3) (t-test, P<0.001).

MS is known to induce a robust CTA (23). Hence, we asked whether rotation could establish a CTA in a two-bottle choice test (37). Two conditioning sessions pairing a 5% sucrose solution with rotational motion, were followed by a choice test of water or sucrose two days later. We observed a significant reduction in sucrose preference when sucrose was paired with rotation, compared to controls (Figure 1E).

With the validity of the rotation paradigm established, we set out to define the neuronal substrate of vestibular-induced, MS-like responses. Excitatory neurons are highly abundant in the VN (30) and glutamatergic VN neurons are activated after provocative motion (35). Among the different glutamatergic markers, VGLUT2 (encoded by *Slc17a6*) shows a robust and significant expression in the VN (38), and VGLUT2-expressing neurons have been shown to participate in vestibular responses to hypergravity and postural imbalance, in contrast to GABAergic (VGAT-expressing) inhibitory neurons (31, 32). Thus, we assessed the necessity of genetically defined glutamatergic VGLUT2-expressing VN (VGLUT2^VN^) neurons in eliciting MS behavioral and autonomic responses by bilateral chemogenetic inhibition in a well-characterized *Slc17a6^Cre^* mouse line (39) (Figure 1F, Figure S1B). Targeted chemogenetic inhibition of VGLUT2^VN^ neurons by expression of hM4Di and administration of clozapine N-oxide (CNO) little effect on locomotion prior to spin stimulation (Figure S1C). However, inhibition of VGLUT2^VN^ neurons by CNO administration prevented spin-induced decreases in ambulatory activity (Figure 1G), appetite suppression (Figure 1H) and attenuated the MS-related decrease in body temperature (Figure 1I). An initial decrease in body temperature was observed in CNO-injected mice, suggesting that other neuronal populations may contribute to the early spin-induced drop in body temperature. In addition, inhibition of VGLUT2^VN^ neurons also prevented spin-induced CTA (Figure 1J). In contrast, bilateral chemogenetic inhibition of GABAergic (*Gad2* -expressing) VN neurons did not alter spin-induced hypolocomotion (Figure S1D-F). These results indicate that VGLUT2^VN^ neurons are necessary to promote the development of the MS-like behavioral and autonomic responses elicited by rotational stimulation.

Nauseogenic responses can be obtained after unilateral inner ear caloric stimulation (40). Hence, to test whether VGLUT2^VN^ neuronal activation is sufficient to induce MS-like autonomic responses, *Slc17a6^Cre^* mice received a unilateral injection of an AAV1 expressing Cre-dependent ChR2:YFP (VGLUT2^VN^:ChR2 mice; for photoactivation) or a YFP construct (VGLUT2^VN^:YFP mice; as control) into the right VN, and an optical fiber was implanted over the injected VN (Figure 2A). Increased photostimulation frequencies ranging from 10 to 40 Hz, reduced ambulatory activity in VGLUT2^VN^:ChR2 mice (Figure S1B). We used a 5-min, 40-Hz, optogenetic-stimulation paradigm because we and others have shown VGLUT2^VN^ neurons in actively moving mice maintain firing rates of up to 40 Hz (41, 42). VGLUT2^VN^ neuron optogenetic activation led to a significant decrease in spontaneous ambulatory activity (Figure 2B), consistent with the results obtained after rotational stimulus. Pairing 5% sucrose solution to unilateral optogenetic activation of VGLUT2^VN^ neurons did not result in significant differences between the ChR2 and the YFP group (Figure 2C), even though ingestion of their regular, low-calorie chow and water intake were significantly decreased after optogenetic VGLUT2^VN^ activation, showing complete suppression for as long as 30 min after laser onset (Figures 2D and 2E). Noteworthy, a normal feeding pattern was observed after presentation of highly palatable, chocolate-flavored, liquid diet to a separate cohort of laser-stimulated VGLUT2^VN^:ChR2 mice (Figure 2F), ruling out physical inability to feed but rather a lack of motivational drive to consume regular chow or water. Core body temperature significantly decreased, with a maximum 3°C drop occurring 17.5 min after laser onset (Figure 2G). To assess whether the observed optogenetic-induced loss of core body temperature was due to a reduction in ambulatory activity in VGLUT2^VN^:ChR2 mice, an additional optogenetic stimulation was applied under physical restraint. Under these conditions, core body temperature increased in VGLUT2^VN^:YFP mice, likely due to restraint-induced stress responses. However, VGLUT2^VN^:ChR2 mice still showed a significant drop in body temperature coincident with photostimulation (Figure 2H). Thus, these results highlight that VGLUT2^VN^ neuron activation is sufficient to recapitulate most MS-induced symptoms.

**Figure 2.**
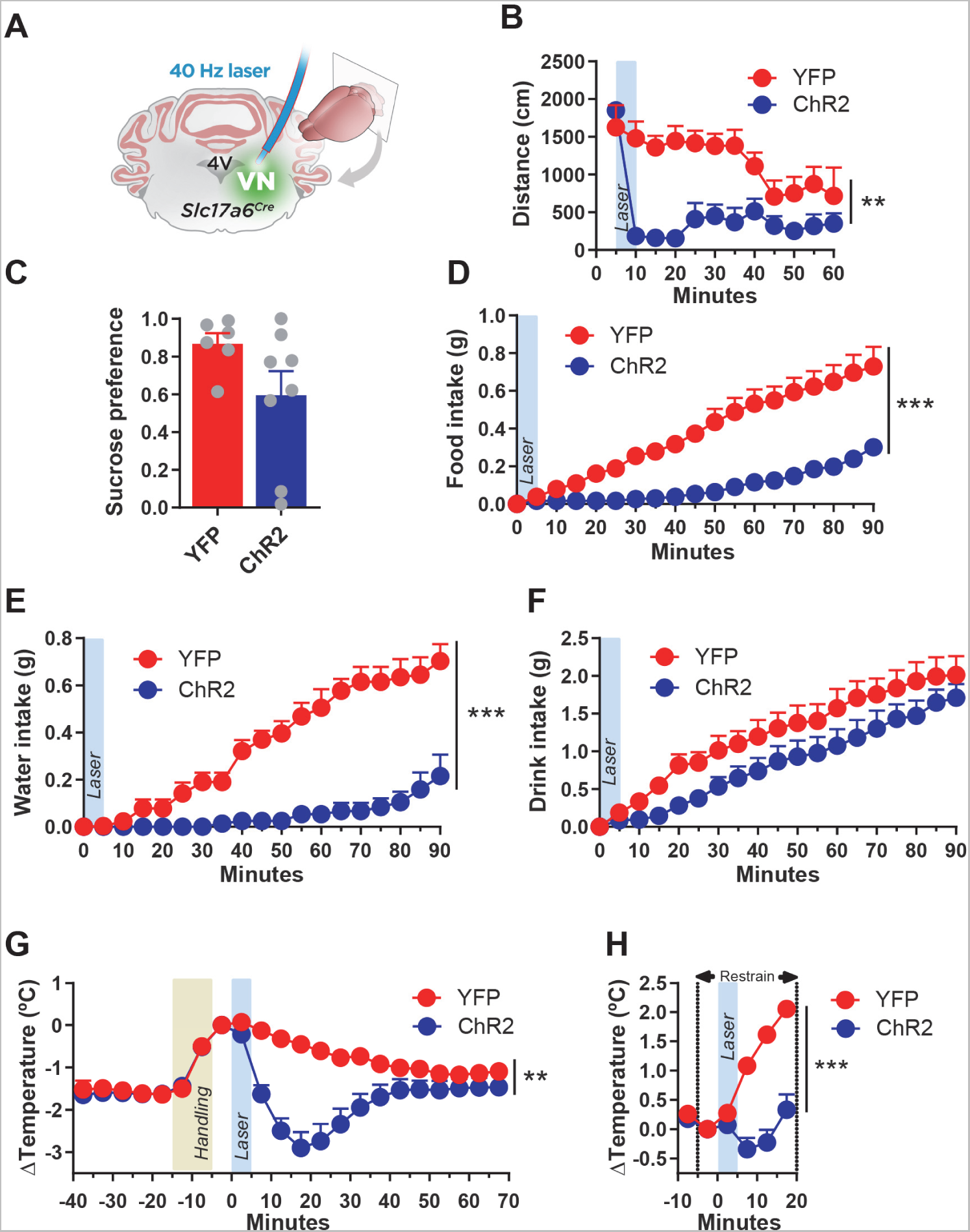
VGLUT2^VN^ neuron activation is sufficient to induce kinetosis. (A) *Slc17a6^Cre^* mice were unilaterally injected in the right VN with AAV1-DIO-ChR2ꞏYFP (VGLUT2^VN^:ChR2 mice) or AAV1-DIO-YFP (VGLUT2^VN^:YFP mice), followed by optical fiber implantation over the right VN to deliver 40-Hz, 10-ms, 10-mW, 473-nm light pulses for 5 min under different behavioral approaches. (B) Traveled distance during 60 min of open-field test after photostimulation (VGLUT2^VN^:ChR2 mice n=7; VGLUT2^VN^:YFP mice n=5; two-way ANOVA, *P*<0.01 effect of ChR2). (C) Conditioned taste avoidance (CTA) response in mice exposed to a two-bottle-based test pairing a 5% sucrose solution to optogenetic activation of VGLUT2^VN^ neurons in VGLUT2^VN^:ChR2ꞏYFP (n=8) or control VGLUT2^VN^:YFP (n=6) mice (t-test, P>0.05). (D) Normal-chow intake following laser onset in VGLUT2^VN^:ChR2 or VGLUT2^VN^:YFP mice (n=6; two-way ANOVA, *P*<0.001 effect of ChR2). Animals were food-deprived for 24 h prior to photostimulation. (E) Water intake following laser onset in food-deprived, VGLUT2^VN^:ChR2 or VGLUT2^VN^:YFP mice (n=6; Two-way ANOVA, P<0.001 effect of ChR2). (F) Highly palatable, chocolate-flavored drink intake following laser onset in food-deprived VGLUT2^VN^:ChR2 or VGLUT2^VN^:YFP animals (n=6; Two-way ANOVA, P>0.05 effect of ChR2). (G) Core body temperature difference (ΔT) in freely moving VGLUT2^VN^:ChR2 or VGLUT2^VN^:YFP mice after handling and laser stimulation (n=8; two-way ANOVA, *P*<0.01 effect of ChR2). (H) Core body temperature difference (ΔT) after photostimulation in restrained VGLUT2^VN^:ChR2 or VGLUT2^VN^:YFP mice (n=5; two-way ANOVA, *P*<0.001 effect of ChR2). YFP: VGLUT2^VN^:YFP mice; ChR2: VGLUT2^VN^:ChR2 mice.

### Identification of *Crh*- and *Cck*-expressing VGLUT2^VN^ neuronal subpopulations

To define VGLUT2^VN^-neuron subpopulations, we performed viral vector-mediated RiboTag molecular profiling (43–45) in VN homogenates of *Slc17a6^Cre^* mice injected with a RiboTag-expressing construct (AAV1-DIO-Rpl22:HA (44); Figure 3A). Transcripts enriched in VGLUT2^VN^ neurons were identified by differential expression analysis in RNA samples extracted from the RiboTag immunoprecipitates (IP; containing polysome-associated mRNAs from VGLUT2^VN^ neurons) and the input (I) of the immunoprecipitation (containing RNA from all the different cell types in the VN). Data analysis confirmed specific enrichment for *Slc17a6* (Vglut2), and depletion of inhibitory neuron mRNAs (*Gad2*) and non-neuronal transcripts (*Cnp*, *Gfap*) in the RiboTag IPs. In addition, there was a significant enrichment for candidate VGLUT2^VN^ neuron subpopulation markers such as *Cck*, *Crh*, *Adcyap1*, *Gal*, *Cbln1* and *Coch* (Figure 3B).

**Figure 3.**
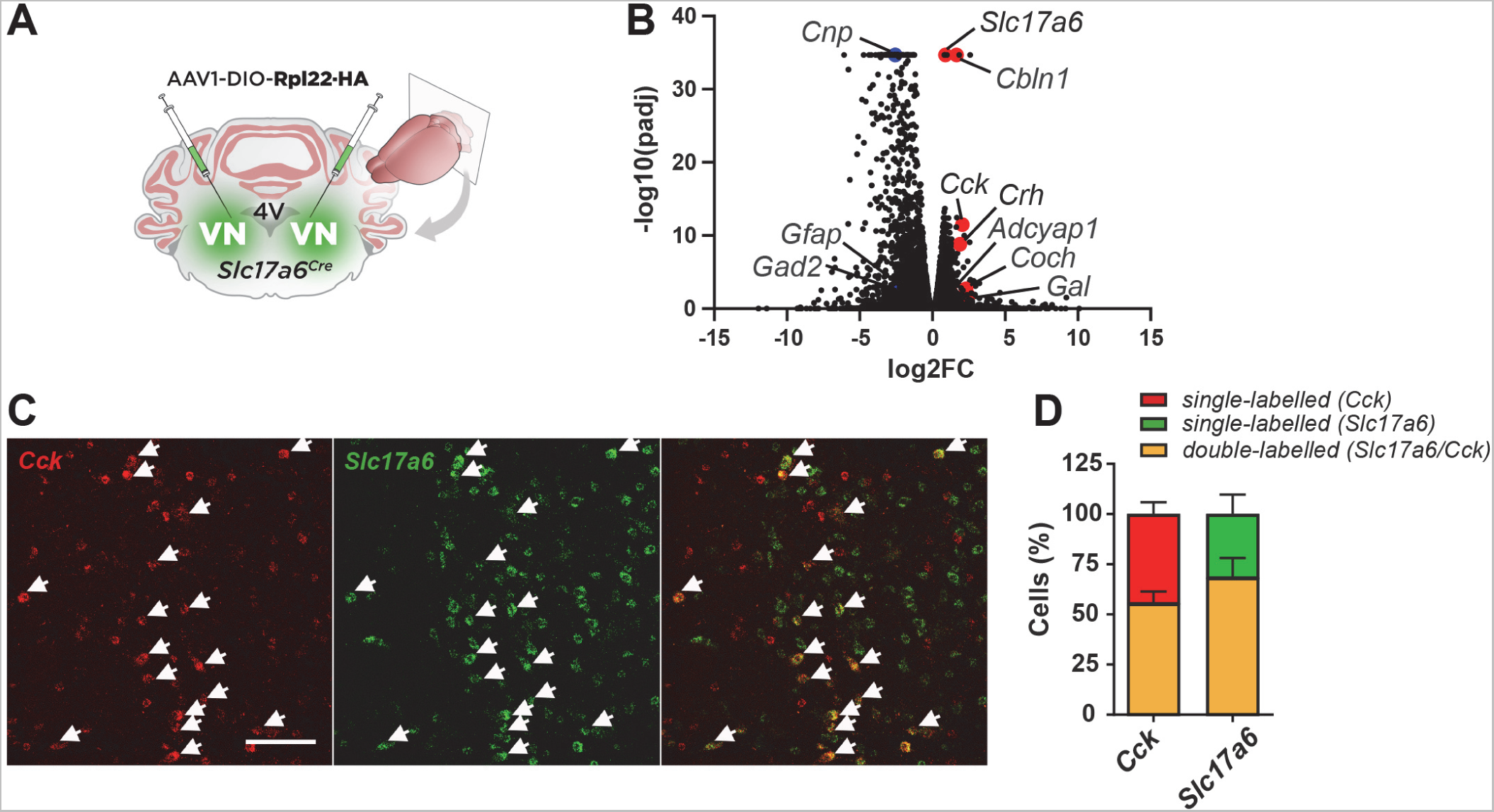
Identification of glutamatergic neuronal subpopulations in the VN. (A) *Slc17a6^Cre^* mice were bilaterally injected in the VN with a viral vector expressing the RiboTag (AAV1-DIO-Rpl22HA) for the molecular profiling of VGLUT2^VN^ neurons. (B) Differential expression analysis showing significant enrichment for candidate VGLUT2^VN^ neuron subpopulation marker transcripts in the immunoprecipitates of RiboTag assays when compared to inputs (n=3). Specific enrichment for *Slc17a6* (Vglut2), and depletion for inhibitory neuron (*Gad2*) and non-neuronal marker transcripts (*Cnp*, *Gfap*) were confirmed in the analysis (padj: adjusted *P*-value; FC: fold change). (C) Double-label *in situ* hybridization assay showing expression of *Cck* mRNA within *Slc17a6-*expressing cells. Scale bar: 100 µm. (D) Percentage of double labelled *Cck*- and *Slc17a6*-expressing cells.

Among these, *Crh*- and *Cck-*expressing neurons have been shown to be involved in autonomic and nauseogenic responses (46, 47), highlighting the potential relevance of these neuronal populations in VN-mediated MS responses. Subsequent *in situ* hybridization assays confirmed the existence of scattered *Crh*-positive cells that co-localized with *Slc17a6* (Figure S2A) and an abundant population of *Cck-*expressing neurons, constituting >60% of all *Slc17a6*- expressing neurons (Figure 3C,D). Noteworthy, a second population of *Cck-*expressing cells (∼45% of the total) that co-localized with *Gad2* was observed (Figure S2B).

### Activation of vestibular *Cck*-expressing neurons causes MS-like autonomic responses

To test the necessity of vestibular *Crh*- or *Cck*-expressing neuronal populations (CRH^VN^ or CCK^VN^ neurons, respectively) in the development of MS-like signs, we injected *Crh^Cre^*or *Cck^Cre^* mice bilaterally in the VN with an AAV vector carrying Cre-dependent hM4Di-mCherry (Figure S3A and 4A). Chemogenetic inhibition of CRH^VN^ neurons did not induce any effect pre- or post-spin on ambulatory activity (Figure S3B,C), indicating that they are not necessary to develop rotation-induced autonomic responses. Unexpectedly, bilateral chemogenetic inhibition of CCK^VN^ neurons significantly decreased ambulatory activity in the absence of rotational stimulus (Figure 4B), which led to an inability to move after the spin (Figure 4C). Reducing the volume of the inhibitory chemogenetic vector (from 0.35 µl to 0.2 µl) in the VN of *Cck^Cre^* mice also led to a marked decrease in locomotion in the CNO-injected hM4Di group, ruling out a role for *Cck*-expressing neurons from neighboring nuclei (Figure S3D-F). Since CCK^VN^ neuron inhibition *per se* is sufficient to induce locomotor effects, further experiments using this approach were hindered. Thus, we sought to establish if stimulation of CCK^VN^ neurons would be sufficient to elicit MS-like responses. Optogenetic activation of CCK^VN^ neurons (Figure 4D), led to a significant and prolonged decrease in spontaneous ambulatory activity in the open-field test compared to the control animals (Figure 4E) and the appearance of a robust CTA (Figure 4F). Similarly, food intake was significantly decreased after photostimulation of CCK^VN^ neurons, showing a complete suppression for 35 min, while control animals engaged in feeding almost immediately (Figure 4G). Optogenetic stimulation of CCK^VN^ neurons reduced core temperature (∼4.5°C), occurring 22.5 min after laser onset, in contrast to control mice (Figure 4H). Thus, our results underscore that stimulation of CCK^VN^ neurons is sufficient to elicit MS-like behavioral and autonomic responses.

**Figure 4.**
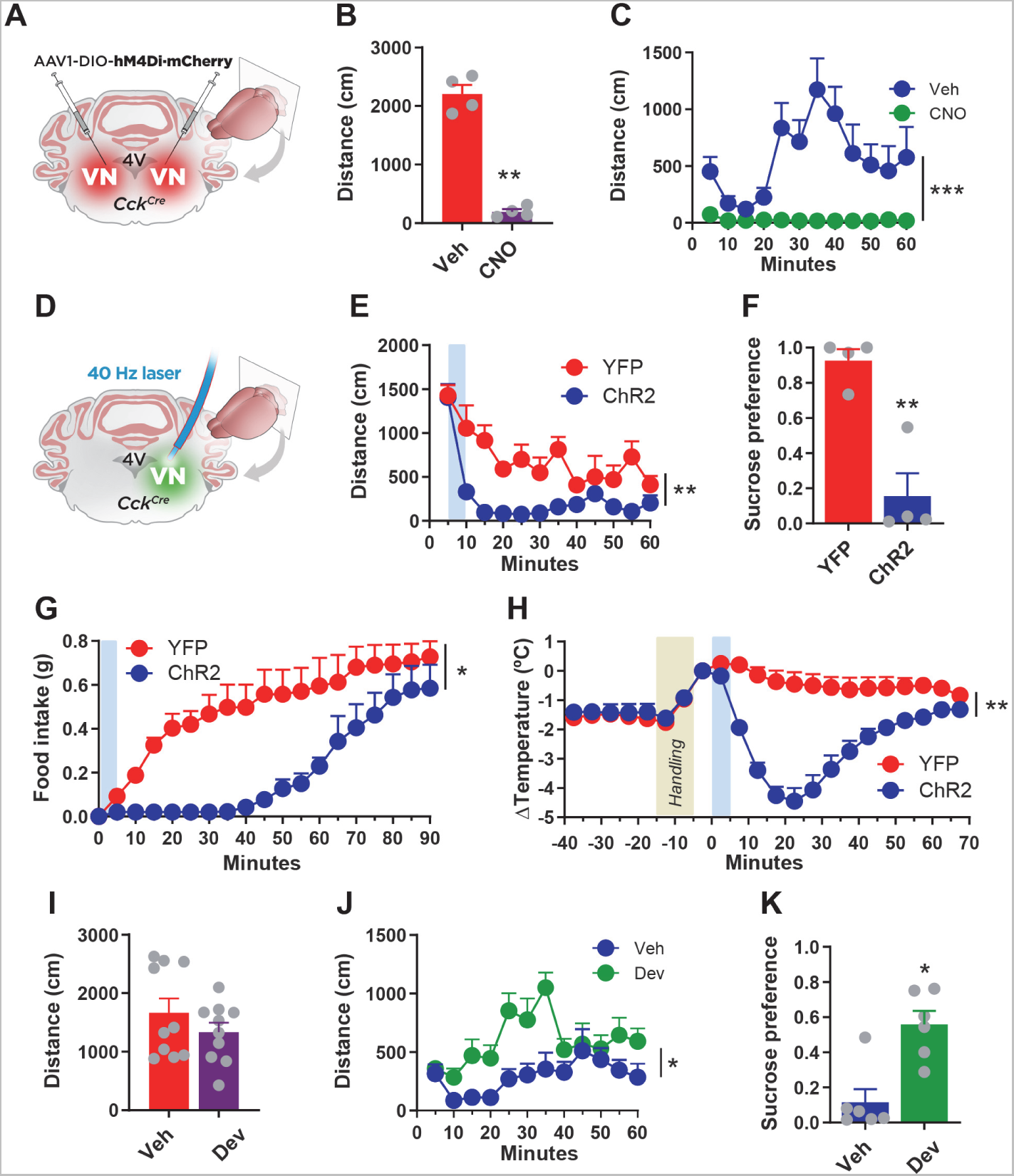
CCK^VN^ neuron manipulation affects motion sickness responses. (A) *Cck^Cre^* mice were bilaterally injected in the VN with AAV1-DIO-hM4DiꞏmCherry (CCK^VN^:hM4Di mice) to inhibit CCK^VN^ neurons upon CNO administration. (B) 5-min open-field test in CCK^VN^:hM4Di mice 30 minutes after CNO or vehicle administration (n=4; t-test, *P*<0.01). (C) Open-field test in CCK^VN^:hM4Di mice injected with CNO or vehicle after spin stimulation (n=4; two-way ANOVA, *P*<0.001 effect of drug). (D) *CCK^Cre^* mice were unilaterally injected in the right VN with AAV1-DIO-ChR2ꞏYFP (CCK^VN^:ChR2 mice) or AAV1-DIO-YFP (CCK^VN^:YFP mice) followed by an optical fiber implantation to deliver a 5-min, 473-nm laser stimulation (40-Hz, 10-mW, 10-ms pulses). (E) Open-field test showing traveled distance after photostimulation in CCK^VN^:ChR2 and CCK^VN^:YFP mice (n=4; two-way ANOVA, *P*<0.01 effect of ChR2). (F) Conditioned taste avoidance (CTA) response in mice exposed to a two-bottle-based test pairing a 5% sucrose solution to optogenetic activation of CCK^VN^ neurons in CCK^VN^:ChR2ꞏYFP or control (CCK^VN^:YFP) mice (n=4; t-test, *P*<0.01). (G) Normal-chow intake after light stimulation in CCK^VN^:ChR2 and CCK^VN^:YFP mice. Mice were food deprived for 24 h prior to stimulation (n=4; two-way ANOVA, *P*<0.05 effect of ChR2). (H) Core body temperature difference (ΔT) in CCK^VN^:ChR2 and CCK^VN^:YFP mice after handling and laser stimulation (n=4; two-way ANOVA, *P*<0.01 effect of ChR2). (I) Spontaneous ambulatory activity of mice 45 min after devazepide or vehicle administration in a 5-min, open-field test (n=10; t-test, *P*>0.05). (J) Traveled distance during 60 min of open-field test after spin stimulation in devazepide or vehicle-treated mice (n=10; two-way ANOVA, *P* <0.05 effect of drug). (K) Conditioned taste avoidance (CTA) response in devazepide or vehicle-treated mice exposed to a two-bottle-based test pairing a 5% sucrose solution to rotational stimulus (n=6; t-test, P-value<0.05).

Since global CCK^VN^ inactivation alters normal VN function, likely by the combined action of neurotransmitters and neuropeptides, we also assessed the specific role of CCK signaling blockade in spin-induced responses (Figure 4I-K). Systemic administration of devazepide (Dev), a CCK-A receptor antagonist, prior to the rotation stimulus, was sufficient to block hypolocomotion (Figure 4J) and the appearance of CTA (Figure 4K), without having any overt behavioral pre-spin effects (Figure 4I) other than an increase in food intake, as described (48, 49) (Figure S3G). In comparison, administration of dimenhydrinate (DMH), a the common anti-MS antihistamine, was able to ameliorate spin-induced hypolocomotion only at 40 mg/kg, a dose that reduced locomotion prior to the rotatory stimulus (Figure S3H,I). These results reveal that vestibular CCK signaling can mediate MS-like behavioral and autonomic responses.

### A CCK^VN^**^-+,^**^PBN^ circuit mediates MS-like CTA via CGRP neuron activation

We hypothesized that specific vestibular projections were likely driving discrete MS-induced responses. To elucidate the genetically defined, vestibular outputs involved in these responses, we compared the vestibular projections described in the Allen Mouse Brain Connectivity Atlas (121146455, 300687607) to the projection fields of *Slc17a6^Cre^*, *Cck^Cre^*, and *Gad2^Cre^* mice injected with a Cre-dependent hM4Di·mCherry construct (Table S1). Among the different brain regions, the PBN was consistently targeted in all animal groups (Figure 5A), in agreement with the Allen Mouse Brain Connectivity Atlas and prior reports (27–29, 35, 50, 51). Furthermore, among brain areas showing robust CCK^VN^ axonal projections, both the VN and the PBN, but not the GRN, had more Fos-positive neurons after the rotational stimulus as assessed by immunohistochemistry (Figure 5B and Figure S4A). Thus, these data confirmed the PBN as a prominent terminal field for VN neurons.

**Figure 5.**
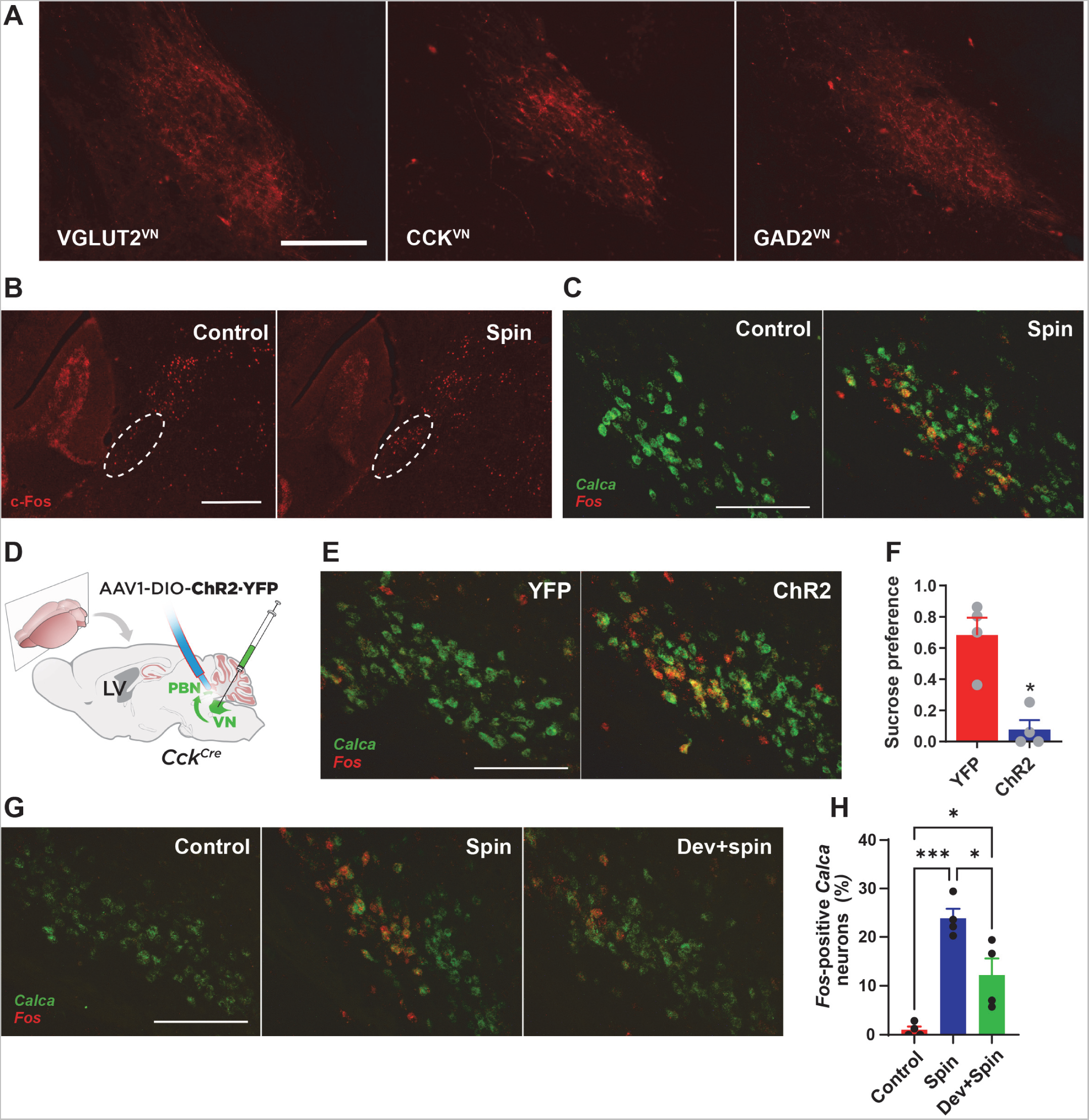
CCK^VN^ neurons target the PBN to mediate CTA. (A) Representative images at Bregma −5.2 mm showing projections to the PBN from VGLUT2^VN^, CCK^VN^ and GAD2^VN^ neurons as assessed by hM4DiꞏmCherry visualization in sections containing the PBN from VGLUT2^VN^:hM4Di, CCK^VN^:hM4Di and GAD2^VN^:hM4Di mice. Scale bar: 150 µm. (B) c-Fos staining at Bregma −5.2 mm from mice subjected to spin or control stimulation. Dotted line delineates lateral PBN. Scale bar: 400 µm. (C) Double-label *in situ* hybridization assay (RNAscope) showing expression of *Fos* mRNA within *Calca-*expressing neurons in the PBN after rotational stimulation. Scale bar: 250 µm. (D) Illustration showing AAV1-DIO-ChR2ꞏYFP or AAV1-DIO-YFP injection in the VN and optical fiber implantation over the PBN in *Cck^Cre^* mice for photoactivation of CCK^VN^ neuron projections in the PBN (CCK^VN◊PBN^). (E) Double-label *in situ* hybridization assay (RNAscope) showing expression of *Fos* mRNA within *Calca-*expressing neurons in the PBN after photoactivation of CCK^VN◊PBN^ projections. Scale bar: 250 µm. (F) CTA response in mice exposed to a two-bottle-based test pairing a 5% sucrose solution to targeted optogenetic activation of the CCK^VN◊PBN^ circuit in CCK^VN^:ChR2ꞏYFP (n=3) or control (CCK^VN^:YFP; n=4) mice (t-test, *P*<0.05). (G) Double-label *in situ* hybridization assay (RNAscope) showing expression of *Fos* mRNA within *Calca-*expressing neurons in the PBN of control or spin-stimulated mice with or without devazepide (Dev; 1 mg/kg) administration. Scale bar: 250 µm. (H) Quantification of the percentage of *Fos-*positive *Calca*-expressing cells in the PBN of control or spin-stimulated mice with or without devazepide (Dev; 1 mg/kg) administration (n=4; one-way ANOVA,* *P* <0.05, *** *P* <0.001).

Our results pointed at an excitatory role for vestibuloparabrachial inputs. Accordingly, to identify the neuronal population in the PBN receiving inputs from VN neurons, we performed double-label ISH assays for *Fos* and *Calca* (which encodes the calcitonin gene-related peptide, CGRP) because these PBN neurons have been implicated in mediating visceral malaise and CTA (37, 52, 53). Results showed that spin stimulation induced *Fos* expression in *Calca*-expressing neurons (Figure 5C). Thus, we hypothesized that a *Cck*-expressing vestibuloparabrachial (CCK^VN➔PBN^) circuit might be relevant in MS by impinging onto CGRP^PBN^ neurons. To that end, we optogenetically activated CCK^VN^ axon terminals in the ipsilateral PBN by placing the optical fiber-tip over this region in CCK^VN^:ChR2 mice (Figure 5D and S5A). Photoactivation of CCK^VN^ neuronal terminals in the PBN increased the number of *Fos* positive-cells in the PBN, which colocalized with *Calca*-expressing neurons (Figure 5E).

Behaviorally, optogenetic stimulation of CCK^VN➔PBN^ fibers at 40 Hz significantly decreased sucrose preference when compared to controls (Figure 5F), as well as body temperature (Figure S5B) without affecting locomotion or food intake (Figure S5C,D). Overall, these results underscore CCK^VN→PBN^ activation of CGRP neurons as a key component of MS-induced CTA. Quantification of activated CGRP^PBN^ neurons after rotational stimulus, as assessed by double ISH assays for *Fos* and *Calca* transcripts (Figure 5G,H) showed that approximately 25% of CGRP^PBN^ neurons respond to spin. Antagonism of CCK-A receptor by devazepide administration led to a 50% decrease in the percentage of active CGRP^PBN^ neurons after rotational stimulus but did not significantly alter the number of *Fos* expressing neurons in the VN (Figure S4B,C). Thus, our results identify a role for CCK-A signaling in the PBN for the development of MS responses.

## DISCUSSION

Being described by Hippocrates over 2000 years ago, MS affects millions of individuals (3). The CNS is thought to compute an MS-triggering sensory conflict signal analogously to “toxic shock”, eliciting malaise and nausea (6, 12, 25, 54). Thus, in an evolutionary context, MS responses could represent a proxy for toxicity-induced balance mismatches. Hence, vomiting observed in humans would be a response to evacuate toxic substances, appetite suppression to avoid additional toxic ingestion, hypolocomotion and hypothermia to minimize the metabolism, and CTA to avoid consumption of toxins in the future. However, the neurobiological underpinnings of MS have remained elusive.

Our results link VGLUT2^VN^ and CCK^VN^ neuronal populations to the development of MS-like symptoms in mice. Furthermore, we reveal a CCK^VN→PBN^ projection that is sufficient to induce CTA and hypothermia likely through the activation of an ensemble of PBN neurons that express the CCK-A receptor including *Calca*-expressing neurons. Our results suggest the potential use of CCK-A receptor blockers as a treatment for MS.

MS is conserved among animal phyla (4, 6–9), but with a high degree of heterogeneity in its behavioral and physiological correlates. Mice do not present an emetic reflex but do present overt physiological and behavioral alterations when exposed to a nauseogenic experience (4, 17, 18, 23, 24, 55) such as sustained 2-*g* rotational stimuli (8, 16–18, 24, 31). We show that an intermittent rotational stimulus of 4 min is sufficient to develop robust hypolocomotion, hypophagia, hypothermia, and CTA. The role of the VN in the development of MS is well recognized (16, 17). The currently accepted hypothesis is that the vestibular inner ear organs provide a major input for the subsequent computing comparisons between present sensory input (integrated input including vestibular, visual and proprioceptive information) and memory recalled from similar motion situations experienced in the past (12, 13). Rotational stimuli convey information from the semicircular canals mostly onto medial VN (MVN) neurons, leading to nauseogenic responses (5, 40, 56). Previous studies had identified that VGLUT2^VN^ neurons, highly abundant in the VN (30), participate in postural balance and mediate gravitational stress-induced hypolocomotion, hypophagia, and hypothermia (31, 32). Here, we confirm these results and show, for the first time, that VGLUT2^VN^ neurons are necessary for the development of MS responses, such as CTA, in a rotational paradigm in mice. Our results show that while unilateral optogenetic activation of VGLUT2^VN^ neurons elicits the responses, bilateral chemogenetic inhibition of the same populations does not have the converse effect on locomotion, which may suggest that a glutamatergic drive is necessary to elicit vestibular-induced effects on locomotion. Alternatively, bilateral inhibition may result in the absence of conflicting commissural inhibition, thus limiting the effects on locomotion (49).

Several excitatory populations have been described in the VN that may be involved in different aspects of vestibular-mediated responses (57). Using the RiboTag approach (43, 45) we identified several genetic markers for subpopulations. Our results validate genes *Crh* and *Adcyap1* (57) and identify new markers for VGLUT2^VN^ neurons including *Gal*, *Coch* and *Cck*. Furthermore, we provide evidence that *Cck*-expressing neurons are the most abundant glutamatergic vestibular neuronal population.

At the functional level, we show that unilateral optogenetic activation of CCK^VN^ neurons is sufficient to recapitulate MS-induced responses, such as hypolocomotion, hypothermia, hypophagia and to elicit a robust CTA. On the other hand, restricted chemogenetic inhibition of CCK^VN^ neurons significantly decreased ambulatory activity and temperature, even in the absence of a rotational stimulus. Different mechanisms may account for this paradoxical response. First, VN presents a high degree of compensation, such as contralateral commissural inhibition (58). Thus, alterations in neuronal activity may lead to compensatory contralateral activation. However, these responses were neither observed after VGLUT2^VN^, GAD2^VN^ nor CRH^VN^ inhibition, pointing at a specific role of CCK^VN^ neurons. In this regard, we have uncovered the existence of both glutamatergic, and GABAergic CCK^VN^ neurons. This fact, given the existence of GABAergic local and commissural interneurons as well as GABAergic projection populations (59) may underlie the strong effect observed after CCK^VN^ activity modulation. Alternatively, MVN neurons produce endogenous, spontaneous pacemaker activity (60). Restoration of MVN pacemaker activity is key for MS habituation (61). Thus, it may be possible that CCK ^VN^ neurons contribute to pacemaker activity and alterations in their rhythmic firing rate may lead to MS-like symptoms. Alternatively, it is possible that either activation or inhibition of these neurons affects balance, which may secondarily cause the MS-responses. Future studies assessing the contribution of balance *per se*, are warranted.

At the circuitry level, our results underscore genetically defined, target- and density-specific projections that may underlie their differential contribution to MS-induced responses. Of the different axonal projections, we describe dense projections from CCK^VN^ neurons to the PBN. The PBN is known to mediate malaise, appetite suppression, lethargy, anxiety, thermoregulation, and CTA (26, 55, 62). Chemogenetic inhibition of VGLUT2^VN^ neurons blocked hypothermia and CTA in our MS paradigm. Unilateral optogenetic stimulation of CCK^VN^ neurons or their projections to the PBN produced hypothermia and robust CTA, which is not recapitulated by optogenetic activation of VGLUT2^VN^ neurons, which may suggest the existence of parallel, or opposing, VGLUT2 ^VN^ neuronal populations/pathways. In contrast, activation of VGLUT2^VN^ neurons leads to hypothermia, likely by decreasing sympathetic tone (31). Recent reports revealed that *Pdyn*-expressing PBN neurons (PDYN^PBN^) regulate hypothermia through projections to the pre-optic area (62), involved in temperature regulation (63). Hence, CCK^VN^ activation of PDYN^PBN^ neurons might mediate the MS-induced drop in body temperature.

Rotational stimulus and CCK^VN^ optogenetic stimulation activated CGRP-expressing, glutamatergic neurons in the lateral PBN, which are known to be involved in malaise, and CTA (26), sustaining the establishment of aversive taste memories (37). Thus, we propose that glutamatergic CCK^VN^ input onto CGRP^PBN^ neurons mediates MS-induced CTA. CCK^VN→PBN^ projections do not seem to participate in MS-induced hypophagia and hypoactivity, even though activation of CGRP^PBN^ neurons is involved in appetite suppression and reduced locomotion (26, 64, 65). Thus, it is likely that CCK^VN^ projections to other brain areas may be responsible for these responses. For example, CCK^VN^ neurons also project to the NTS, which contains neurons that can activate PBN^CGRP^ neurons to promote anorexia (66). Alternatively, it is possible that unilateral CCK^VN→PBN^ activation was not sufficient to recruit a large population of CGRP^PBN^ neurons. Accordingly, our results show that ∼25% of PBN^CGRP^ neurons respond to CCK^VN→PBN^ projection activation. Given that different PBN^CGRP^ subpopulations have been recently described (67), this may indicate that only specific subsets of PBN^CGRP^ are targeted by CCK^VN→PBN^ projections. Alternatively, while CTA is commonly referred to as conditioned taste aversion, we acknowledge that we have not assessed other unequivocal signs of aversion (such as gapes and chin rubs) or malaise. Thus, the responses observed may be due to conditioned avoidance, rather than aversion, which are two independent processes (68), which may also explain the absence of a hypophagic response after CCK^VN→PBN^ stimulation.

Our data show that alterations in the activity of CCK^VN^ neurons exert a pivotal role in controlling MS-like behavioral responses. Here we provide several lines of evidence that CCK signaling, through its CCK-A receptor, underlies MS responses. Devazepide administration, abolishes both hypolocomotion and CTA after a rotational stimulus, without affecting spontaneous locomotion like dimenhydrinate, a widely used anti-motion sickness drug.

Pharmacologically, three anti-MS drug classes have been described depending on their influence over MS habituation. Thus, class A drugs such as amphetamine are thought to block the MS-eliciting sensory input, leading to habituation delay. Class B drugs such as anticholinergics modulate the neural store decreasing the neuronal mismatch signal intensity, leading to boosted habituation. On the other hand, class C drugs such as antihistamines inhibit MS-related autonomic responses, leading to unchanged MS habituation (69). Even though we have not tested habituation *per se*, our results suggest that devazepide may be preventing the expression of autonomic responses, akin to class C drugs. Mechanistically, we show that CCK-A receptor blockade reduces CGRP^PBN^ activation, in agreement with our recent study reporting the expression of *Cckra* in CGRP^PBN^ neurons (67). In addition to its known gastric effects, CCK-A receptor signaling has been suggested to contribute to multiple central functions, such as anxiety, nociception, and food consumption (70).

Our study underscores a key role for CCK^VN^ neurons in MS-related behavioral and physiologic responses by impinging onto PBN circuitry, providing the first evidence of a direct link between motion inputs and aversive responses. Furthermore, we identify CCK-A receptor blockade as a novel therapeutic approach for MS. Future studies detailing the behavioral and physiological contribution of other CCK^VN^ targets will provide a more complete profile of the neurobiological substrates of MS.

## ACKNOWLEDGMENTS

Authors thank Diane Durnam for editing the manuscript. This work was supported by a NENS exchange grant for training stay (PMM), a Marie Sklodowska-Curie Individual Fellowship (H2020-MSCA-IF-2014-658352; ES), pre-doctoral fellowships (2018FI_B 00452 to AU; PRE2018-083179 to LSB, PRE2021-096944 to MG) and three Ramón y Cajal fellowships (RyC-2012-11873; AQ, RYC2019-028501-I; ES, RYC2020-029596-I; E.P). E.P received funds from MICINN (PID2021-125079OA-I00). E.S received funds from MICIU Proyectos I+D+i “Retos Investigacion” (RTI2018-101838-J-I00) and MICINN Proyectos I+D+i (PID2019-107633RB-I00 and PID2022-142544OB-I00). A.Q. received funds from the European Research Council (Starting grant NEUROMITO, ERC-2014-StG-638106), MINECO Proyectos I+D de Excelencia (SAF2014-57981P; SAF2017-88108-R), MICINN Proyectos I+D+i (PID2020-114977RB-I00), AGAUR (2017SGR-323, 2021SGR-720), Fundació TV3-La Marató (202030), and “la Caixa” Foundation (ID 100010434), under the agreement LCF/PR/HR20/52400018.

## AUTHOR CONTRIBUTIONS

RP, ES and AQ conceived, designed, and supervised the work. PMM, LSB, FM, AU, MG, EP, IA, ES and AQ acquired, analyzed, and interpreted the data. The manuscript was written by PMM, RP, ES and AQ and revised and edited by all authors.

## METHODS

### Mice

The following mouse lines were used in this study: *Slc17a6^Cre^* (BAC-Vglut2-Cre) (39) mice were generated by Ole Kiehn. *Cck^Cre^* (CCK-IRES-Cre) and *Crh^Cre^* (CRH-IRES-Cre) (71) mice were obtained from The Jackson Laboratory (Bar Harbor, ME. Stock No: 012706 and 012704, respectively). Mice were group-housed with a 12:12h light:dark cycle at 22°C, with *ad libitum* access to rodent chow (Teklad Global Rodent Diet #2014S; Envigo) and water, unless otherwise stated. Sex and age-balanced groups of 2-to 7-month-old mice were used across all experimental procedures. No sex differences were observed. After surgeries, animals were individually housed until the end of all experimental procedures. Sample sizes were determined using power analyses. The number of animals used per group in each experiment (n) are provided in figure legends. All mice were on a C57BL/6J background after backcrossing for at least 10 generations. All experiments were conducted following the recommendations in the Guide for the Care and Use of Laboratory Animals and were approved by the Animal Care and Use Committee of the Universitat Autònoma de Barcelona and the Generalitat de Catalunya.

### Drugs

Devazepide (Dev; 1 mg/kg in saline solution containing 1% DMSO and 1% Tween-80) and dimenhydrinate (DMH; 20 or 40 mg/kg in saline solution) were administered intraperitoneally (i.p.) 45 or 30 minutes before tests, respectively. A between-subject design was used for pharmacologic studies, with animals receiving either Veh-Drug or Drug-Veh in a balanced manner (Figure S6)

### Rotational stimulus

Prior to rotational stimulation, animals were habituated to physical restraint for 4 min using a 50-mL conical tube coupled to a custom-made rotary device (external radius: 10.5 cm; internal radius from mouse head: 5 cm. Rotation multiplier: x3.6). Afterwards, rotational (four repeated 1-min, 4-*g* accelerations) or control (4 min with no rotation) stimuli were applied, unless otherwise stated. 1-min accelerations included 55 s of rotation plus 5 s break until full stop. To achieve 4-*g* accelerations, 75 rpm were applied.

### Behavioral assays

Each animal was subjected to an open-field test, followed by food intake and conditioned taste aversion (CTA) analysis. After tests, telemetric temperature sensors were implanted, and core-body temperature monitored.

### Open-field test

The open-field (OF) test was conducted in a non-covered, white methacrylate box (56 x 36.5 x 31 cm) that allows for video recording during animal testing. Mice were individually exposed for 5 min to the OF before receiving rotational or control stimulation. After stimulation, mice were re-exposed to the OF for 60 more min. For optogenetic experiments, mice were exposed for a total of 60 min to the OF (5 min of pre-stimulation, followed by 5 min of laser stimulation and 50 min of post-stimulation). Spontaneous ambulatory activity was monitored using a video tracking software (Ethovision XT 11.5; Noldus Information Technology).

### Food and liquid intake measurement

Animals were individually placed in an Oxyletpro-Physiocage monitoring system (Panlab) for real-time quantification of food and liquid intake. Mice were habituated to the cage for 2 to 4 days with *ad libitum* access to normal chow and water, unless otherwise stated. Prior to each session, animals were food deprived for 24 h and refed at the onset of the dark cycle. For specific experiments, chocolate-flavored, highly palatable liquid diet (ENSURE Nutrivigor, Abbott) was provided during cage habituation and in the subsequent experimental sessions. Food and liquid intake were expressed as cumulative intake using the Metabolism software version v3.0.00 (Panlab).

### Conditioned taste avoidance (CTA) test

For the CTA test, a two bottle-based protocol was used (37). Animals were individually placed in a custom cage with angular ports for two liquid-containing bottles. *Ad libitum* water access was provided for 2 days during habituation. Next, animals only had access to both water-containing bottles during restricted time periods (30 min access in the morning, 30-60 min in the afternoon) for 8 days (D1-D8). During days 4 and 6 (D4 and D6), a solution of 5% sucrose in water was paired with specific stimuli (rotation, CNO injection, photostimulation) to establish conditioning. Mice were tested on D8. Sucrose preference value was calculated as sucrose solution consumption/total liquid consumption.

### Surgical implantation of telemetry devices and temperature monitoring

Anesthetized mice (5% isoflurane for induction, 1.5% for maintenance) were aseptically implanted with telemetric temperature transmitters (G2 E-Mitter, STARR Life Sciences Corp.) into the peritoneum and allowed to recover for 2 weeks. Mouse cages were placed on telemetry receivers (ER4000 Energizer/Receiver, STARR Life Sciences Corp.) and core body temperature was monitored using the VitalView software version 5.0 (STARR Life Sciences Corp.) under controlled ambient temperature (22°C). All recordings started in resting animals. For specific experiments, mice were inserted into a modified 50-mL Falcon tube to ensure physical restrain while allowing optogenetic tethering.

### Viral vector production

pAAV-hSyn-DIO-hM4Di:mCherry, pAAV-EF1a-DIO-eYFP and pAAV-EF1a-DIO-ChR2:YFP plasmids were obtained from Addgene (#44362, #27056 and #100056, respectively). pAAV RiboTag virus (pAAV-EF1a-DIO-Rpl22·HA) has been described (44). Recombinant adeno-associated viral vectors (AAV) were produced in human embryonic kidney (HEK293T) cells with AAV1 coat serotype. Purification was achieved by several sucrose and CsCl gradient centrifugations and a final re-suspension in 1x Hanks Balanced Saline Solution (HBSS) at a titer of 2 x 10^9^ viral genomes/μL as described (44, 72). AAV preparations were aliquoted and stored at −80°C until stereotaxic injection.

### Stereotaxic surgery

All surgeries were performed under aseptic conditions. Animal anesthesia was induced and maintained with 5% and 1-1.5% isoflurane/O_2_, respectively. Analgesia (5 mg/kg ketoprofen; Sanofi-Aventis) and ocular protective gel (Viscotears®, Bausch+Lomb) were applied. Mice were then placed over a heating pad in a robot-operated, 3-dimensional (stereotaxic) frame (Neurostar) for intracerebral virus delivery. Stereotaxic coordinates were normalized using a correction factor (Bregma-Lambda distance/4.21) based on the coordinates of Paxinos and Franklin(73). AAV preparations were unilaterally (right side) or bilaterally delivered into the VN (antero-posterior (AP), −6.00 mm from Bregma; medio-lateral (ML), ± 0.90 mm; dorso-ventral (DV), −4.00 mm from skull surface) at a constant rate of 0.1 μL/min for 3.5-4.0 min (0.20-0.40 μL per injection site) using a 32-gauge blunt needle coupled to a 5 μL-syringe (Hamilton). After infusion, the needle was maintained in place for 6 min to allow proper diffusion. Subsequent needle withdrawal was performed at 1 mm/min to ensure minimal off-target viral leakage. After viral injection, mice used for optogenetic experiments also received unilateral surgical implantation of a fiber-optic cannula as described below. Only animals with correct targeting were included in the experiment (Figure S7).

### Chemogenetics

*Slc17a6^Cre^, Cck^Cre^* or *Crh^Cre^* mice were bilaterally injected in the VN with 0.35 µL (per side) of AAV1-hSyn-DIO-hM4Di:mCherry. Clozapine-n-oxide (CNO; 1 mg/kg) was administered via intraperitoneal (i.p.) injection 35 min prior to rotational or control stimulation. A between-subject design was used for chemogenetics studies, with animals receiving either Veh-CNO or CNO-Veh in a balanced manner (Figure S6).

### Optogenetics

*Slc17a6^Cre^* or *Cck^Cre^* mice were injected in the right VN with 0.4 μL of either AAV1-EF1α-DIO-ChR2·YFP or AAV1-EF1α-DIO-eYFP and a fiber-optic cannula (200-μm fiber core diameter, 0.22-numeric aperture; 2.5-mm ferrule diameter; Thorlabs) was implanted over the right VN (AP, −6.00 mm; ML, +0.90 mm; DV, −4.00 mm) or over the right PBN (AP, −5.20 mm; ML, +1.70 mm; DV, −3.00 mm) of *Cck^Cre^* animals. A fiber-optic cannula was fixed to the exposed skull with a layer of adhesive cement (Super-Bond C&B, Sun Medical) and dental acrylic cement (Rebaron, GC Corporation). The skin was affixed to the cement with tissue adhesive (Vetbond, 3M). A blue 473-nm laser light was produced by a DPSS Laser System (LRS-0473-GFM-00100-05 Laserglow) and driven by a fiber-optic patch cord (200-μm core diameter, 0.22-numeric aperture: FT030 protection, Thorlabs). Light intensity was set at 10 mW measured by a photometer (Thorlabs) at the tip of a non-implanted fiber optic cannula attached to the patch-cord. To deliver illumination to the right VN or right PBN, the patch cord was connected to the implanted fiber-optic-containing cannula through a ceramic sleeve. A pulse generator (33500B Series Trueform, Keysight) was used to adjust laser output to deliver 40-Hz, 10-ms, pulse trains for 5 min to all mice.

### Mapping of neuronal projections

To visualize the projections of VGLUT2^VN^, CCK^VN^ or GAD2^VN^ neurons, *Slc17a6^Cre^*, *Cck^Cre^* or *Gad2^Cre^* mice were injected into the right VN with 0.2 µL of AAV1-hSyn-DIO-hM4Di:mCherry. Animals were euthanized 3 weeks after surgery for direct visualization of fibers after tissue fixation and cryopreservation.

### Tissue processing and Immunofluorescence analysis

Mouse brains were freshly dissected following euthanasia by CO_2_ asphyxia, fixed overnight with phosphate-buffered saline (PBS) containing 4% paraformaldehyde (PFA) and cryo-protected with 30% sucrose in PBS. For cryo-sectioning, brains were frozen for 5 min in dry ice and sectioned in a freezing microtome. For immunofluorescence, 30-μm free-floating sections were blocked in PBS with 10% normal donkey serum (NDS) and 0.2% Triton X-100 for 1 h at room temperature followed by an overnight incubation at 4°C with primary antibody solution containing a rabbit anti-c-Fos (1:1,000; #ab222699, Abcam) antibody in PBS with 1% NDS and 0.2% Triton X-100. After three washes in PBS with 0.2% Triton X-100, a secondary antibody solution containing a secondary antibody conjugated to Alexa Fluor 594 fluorophore (1:500; Invitrogen) was added to the sections and incubated for 1 h at room temperature. After the incubation, sections were washed three times for 5 min in PBS with 0.2% Triton X-100 and mounted onto slides with DAPI Fluoromount (#17984-24, Electron Microscopy Sciences) before visualization with an EVOS imaging system (Thermo Fisher Sci).

### RiboTag assays

For genetic identification of neuronal subsets, *Slc17a6^Cre^* mice were bilaterally injected in the VN with 0.4 μL of the RiboTag viral vector (AAV1-DIO-Rpl22HA) (44). Animals were euthanized 3 weeks after surgery for subsequent RiboTag analysis. To isolate the polysome-associated mRNAs from VGLUT2^VN^ neurons, punches containing the VN of *Slc17a6^Cre^* mice injected with AAV1-DIO-Rpl22·HA were pooled and homogenized in 1 ml of buffer as described (45). After centrifugation, 4 μL of anti-HA antibody (MMS-101R, 2–3 mg/ml; Covance) was added to 800 μL of the cleared lysate and incubated for 4 h at 4°C. Remaining lysate was saved as input sample. After incubation, 200 μL of protein A/G magnetic beads (Thermo Scientific) were added and incubated overnight at 4°C with rotation. Immunoprecipitates (IPs) were washed in high-salt buffer and RNA from inputs and IPs were extracted(43). For differential expression analysis, 10 ng of RNA was amplified using the Ovation Pico SL WTA system (NuGEN). Fidelity of amplification was confirmed by qPCR analysis of the resulting cDNA using the QuantiTect kit (Qiagen) before biotinylation according to the EncoreIL biotinylation kit (NuGEN). Biotinylated cDNA was quantified, and product size distribution was analyzed using the 2100 Bioanalyzer system with the RNA 6000 Nano chips (Agilent Technologies). Biotinylated cDNA (750 ng) was hybridized at 48°C to MouseRef-8 v2 expression beadchips (Illumina) for 16 h before washing and analyzing according to the manufacturer’s directions. Signal was detected using a BeadArray Reader (Illumina), and data were analyzed for differential expression using the GenomeStudio data analysis software (Illumina). Average normalization, the Illumina custom error model, and multiple testing corrections using the Benjamini-Hochberg false discovery rate were applied to the analysis. Only transcripts with a differential score of >13 (p < 0.05) were considered. Normalized and raw data have been deposited in the National Center for Biotechnology Information Gene Expression Omnibus and are accessible through GEO Series accession number <u>GSE167672</u> .

### In situ hybridization assays

Mouse brains were fresh-frozen in Tissue-Tek O.C.T. compound (Sakura) with dry ice and stored at −80 °C until cryosectioning. Coronal sections (15 µm) containing the VN or the PBN were used for RNAscope (Advanced Cell Diagnostics) analysis following manufacturer’s directions. The following probes were used: Mm-*Slc17a6* (#319171-C3), Mm-*Cck* (#402271-C2), Mm-*Crh* (#316091-C1), Mm-*Gad2* (#439371-C1), Mm-*Fos* (#316921-C1) and Mm-*Calca* (#578771-C2). All *in situ* hybridization assays were imaged using a confocal (Leica SP5) or epifluorescence (Nikon Eclipse 90i) microscope and analyzed in ImageJ (Fiji v1.0) or QuPath open-source software (74). Cell counting was performed in VN sections from Bregma −5.68mm to −6.64mm in 2-3 slices/animal (n=3 animals) with ImageJ Cell Counter plugin (Fiji v1.52). Number of *Fos* positive *Calca*-expressing cells was analyzed using QuPath. Cell segmentation was accomplished in DAPI-stained sections containing *Calca*-expressing neurons from Bregma −5.02 mm to −5.34 mm using the cell detection feature with a cell expansion of 30 µm. After cell detection, *Fos* and *Calca* transcripts were detected using the subcellular detection module. Each section was visually inspected to confirm accurate spot identification. Number of *Fos* positive *Calca*-expressing cells was obtained from 2-5 sections per mouse with 4 mice per group.

### Statistics

Data are shown as the mean ± SEM. GraphPad Prism v9.0 software was used for statistical analyses. Appropriate tests were selected depending on the experimental design as stated in the figure legends. Statistical significance, when reached (p<0.05 was considered significant), is stated in the figure legends. Number of mice in an experiment (n) represents the number of animals per group. Different cohorts of mice were assigned to different tests to avoid undergoing repeated testing. No attrition was observed.

