## Supplemental Figures for "Vestibular CCK signaling drives motion-induced malaise"

### Figure S1

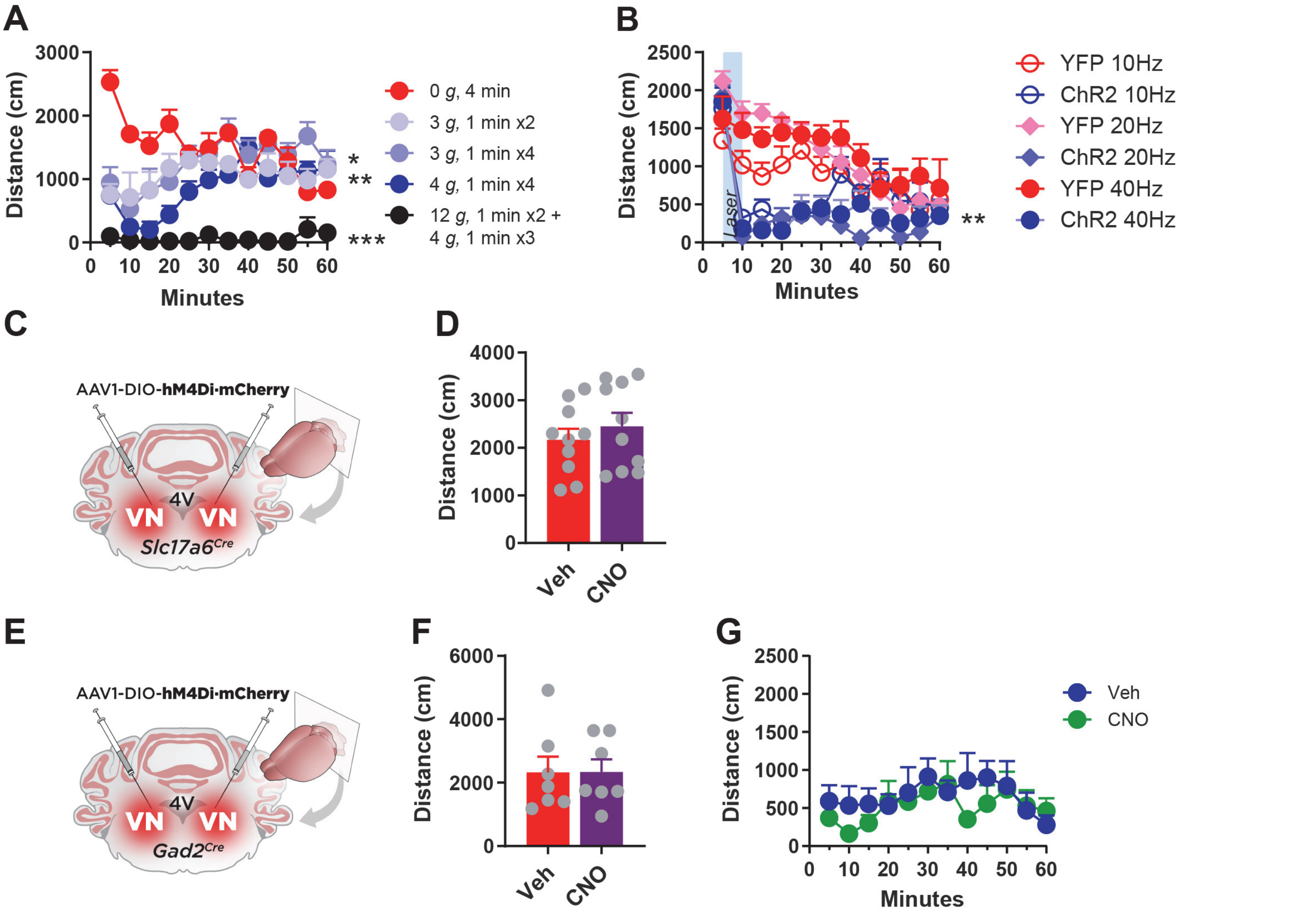


**Figure S1. Effect of increasing rotational intensities, VGLUT2$^{VN}$ optogenetic stimulation, and VGLUT2$^{VN}$ and GAD2$^{VN}$ neuron chemogenetic inhibition on ambulatory activity in mice.** (A) Open-field (OF) test showing reduced locomotor activity in mice subjected to rotational stimuli of different intensity and duration. Centrifugation at 234, 270 and 468 rpm produce a centrifugal force of 3, 4 (n=6) and 12 *g* (n=2), respectively (Two-way ANOVA, *P*<0.001 effect of spin). (B) *Slc17a6$^{Cre}$* mice were unilaterally injected in the vestibular nuclei (VN) with AAV1-DIO-ChR2·YFP (VGLUT2$^{VN}$:ChR2 mice) to activate glutamatergic neurons upon light (473 nm) delivery at 10, 20, or 40 Hz frequencies. Spontaneous ambulatory activity of VGLUT2$^{VN}$:ChR2 mice after 5-min stimulation in an open-field test (n=5-7; two-way ANOVA, *P*<0.01 effect of ChR2 for all stimulation frequencies). (C) *Slc17a6$^{Cre}$* mice were bilaterally injected in the vestibular nuclei (VN) with AAV1-DIO-hM4Di·mCherry (VGLUT2$^{VN}$:hM4Di mice) to inhibit glutamatergic neurons upon CNO administration. (D) Spontaneous ambulatory activity of VGLUT2$^{VN}$:hM4Di mice 30 min after CNO or vehicle administration in a 5-min, open-field test (n=10; t-test, *P*>0.05). (E) *Gad2$^{Cre}$* mice were bilaterally injected in the vestibular nuclei (VN) with AAV1-DIO-hM4Di·mCherry (GAD2$^{VN}$:hM4Di mice) to inhibit GABAergic neurons upon CNO administration. (F) Spontaneous ambulatory activity of GAD2$^{VN}$:hM4Di mice 30 min after CNO or vehicle administration in a 5-min, open-field test (n=7; t-test, *P*>0.05). (G) Total distance traveled during a 60-min open-field session after spin stimulation in CNO- or vehicle-injected GAD2$^{VN}$:hM4Di mice (n=7; Two-way ANOVA, *P*>0.05 effect of drug).

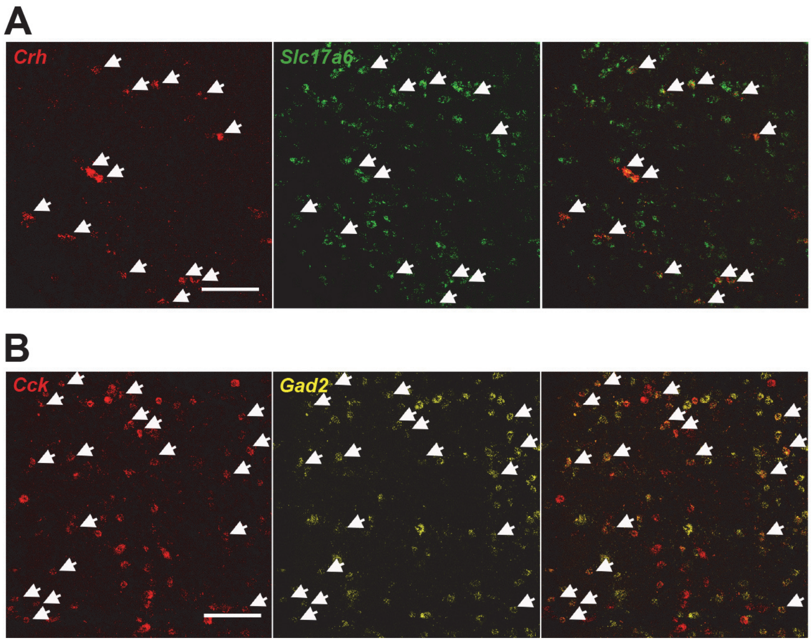


**Figure S2. Excitatory or inhibitory neuron transcript co-expression in CRH$^{VN}$ or CCK$^{VN}$ neurons.** (A) Double-label *in situ* hybridization assay (RNAscope) showing expression of *Slc17a6* mRNA within *Crh*-expressing cells. Scale bar: 100 µm. (B) Double-label *in situ* hybridization assay (RNAscope) showing expression of *Gad2* mRNA within *Cck*-expressing cells. Scale bar: 100 µm.

**Figure S3**

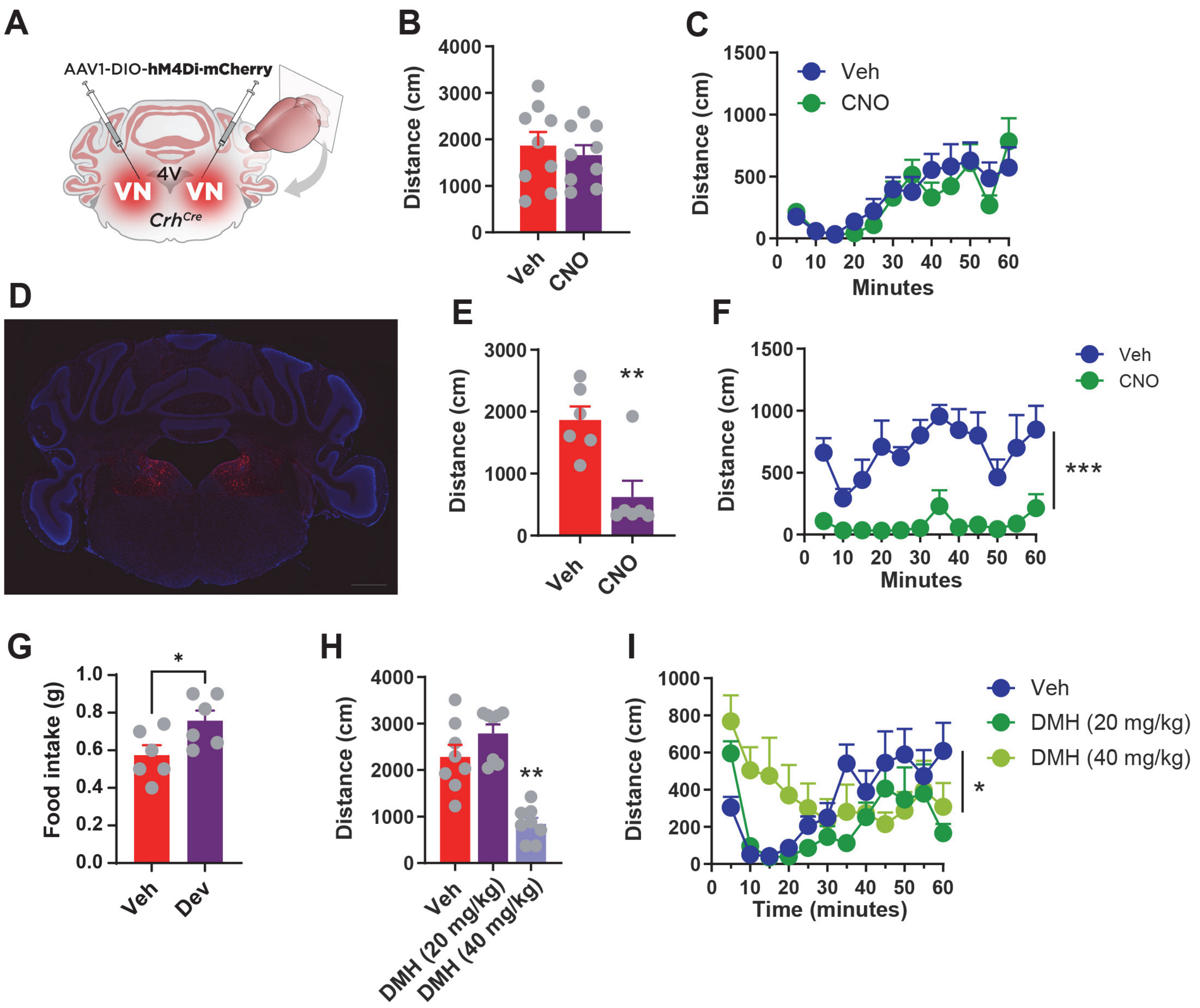


**Figure S3. Effect of chemogenetic inhibition of $CRH^{VN}$ neurons, $CCK^{VN}$ neurons, and devazepide or dimenhydrinate administration on food intake and ambulatory activity before and after spin stimulation.** (A) $Crh^{Cre}$ mice were bilaterally injected in the VN with AAV1-DIO- hM4Di-mCherry ($CRH^{VN}$:hM4Di mice) to inhibit $CRH^{VN}$ neurons upon CNO administration. (B) Locomotor activity in $CRH^{VN}$:hM4Di mice (in a 5-min, open-field session) 30 min after CNO or vehicle administration; (n=9; t-test, $P>0.05$). (C) Locomotor activity in $CRH^{VN}$:hM4Di mice after spin stimulation in CNO and vehicle-injected mice (n=9; Two-way ANOVA, $P>0.05$ effect of drug). (D) Representative image at Bregma -6.0 mm showing restricted hM4Di·mCherry expression in the VN of $CCK^{VN}$:hM4Di mice injected with reduced amounts (0.2 µl) of the viral vector preparation. Scale bar: 800 µm. (E) 5-min open-field test in $CCK^{VN}$:hM4Di mice injected with reduced amounts (0.2 µl) of the viral vector preparation, 30 minutes after CNO or vehicle administration (n=6 ; t-test, $P<0.01$). (F) Open-field test in $CCK^{VN}$:hM4Di mice injected with reduced amounts (0.2 µl) of the viral vector preparation and CNO or vehicle, after spin stimulation (n=6; Two-way ANOVA, $P<0.001$ effect of drug). (G) Cumulative 1-hour food intake of mice 30 min after devazepide (DEV; 1 mg/kg) or vehicle administration (n=6; t.test, $P<0.05$). (H) Spontaneous ambulatory activity of mice 30 min after dimenhydrinate (DMH; 20 mg/kg or 40 mg/kg) or vehicle administration in a 5-min, open-field test (n=8; t-test, $P>0.05$). (I) Traveled distance during 60 min of open-field test after spin stimulation in dimenhydrinate (DMH; 20 mg/kg or 40 mg/kg) or vehicle-treated mice (n=8; Two-way ANOVA, $P<0.05$ effect of drug).

Figure S4

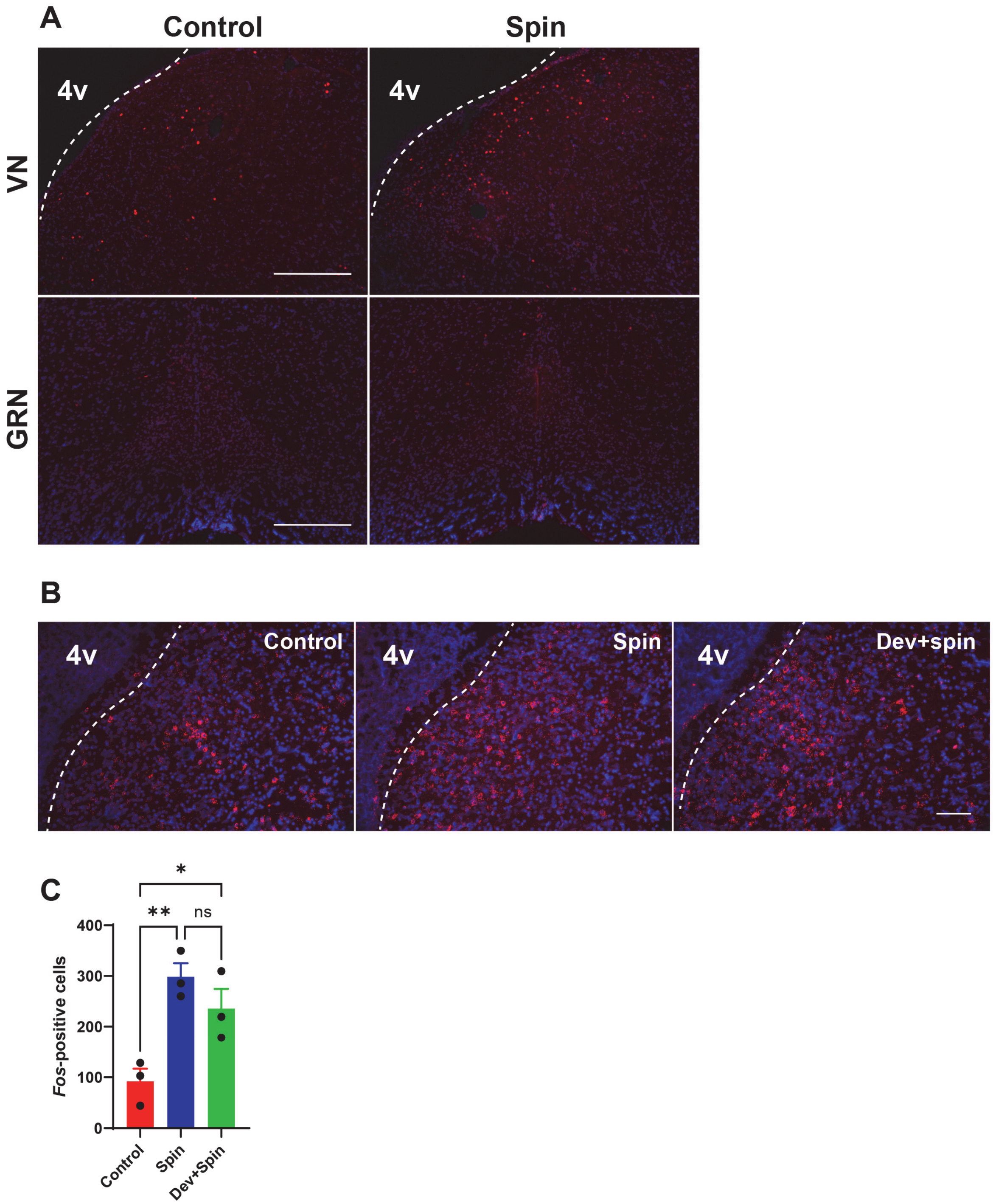


**Figure S4. Effect of rotational stimulus and devazepide administration on neuronal activation in CCK$^{VN}$ projection areas**

(A) Fos staining in the VN and GRN from mice subjected to spin or control stimulation. Dotted line delineates separation between VN and 4th ventricle (4v). Scale bar: 300 µm. (B) Double-label *in situ* hybridization assay showing expression of *Fos* mRNA in the VN of control or spin-stimulated mice with or without devazepide (Dev; 1 mg/kg) administration. Scale bar: 100 µm. (C) Quantification of the percentage of *Fos*-positive cells in the VN of control or spin-stimulated mice with or without devazepide (Dev; 1 mg/kg) administration (n=3).

**Figure S5**

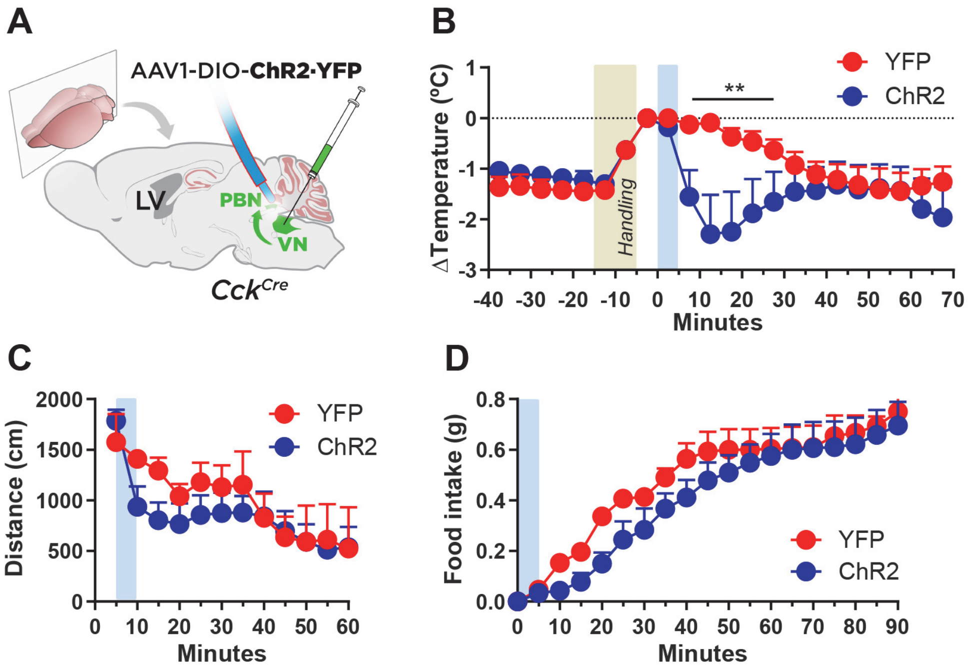


**Figure S5. Effect of the photoactivation of the CCK$^{VN→PBN}$ projection.** (A) Illustration showing AAV1-DIO-ChR2·YFP or AAV1-DIO·YFP injection in the VN and optical fiber implantation over the PBN in *Cck$^{Cre}$* mice for photoactivation of CCK$^{VN}$ neuron projections in the PBN (CCK$^{VN→PBN}$). (B) Core body temperature difference (ΔT) after handling and CCK$^{VN→PBN}$ optogenetic stimulation (CCK$^{VN→PBN}$:ChR2 mice n=3; CCK$^{VN→PBN}$:YFP mice n=4; Two-way ANOVA, $P$<0.01 effect of ChR2). (C) Open-field test showing traveled distance after CCK$^{VN→PBN}$ photostimulation (n=4; Two-way ANOVA, $P$>0.05 effect of ChR2). (D) Food intake in food-deprived mice after CCK$^{VN→PBN}$ photostimulation (n=3; two-way ANOVA, $P$>0.05 effect of ChR2).

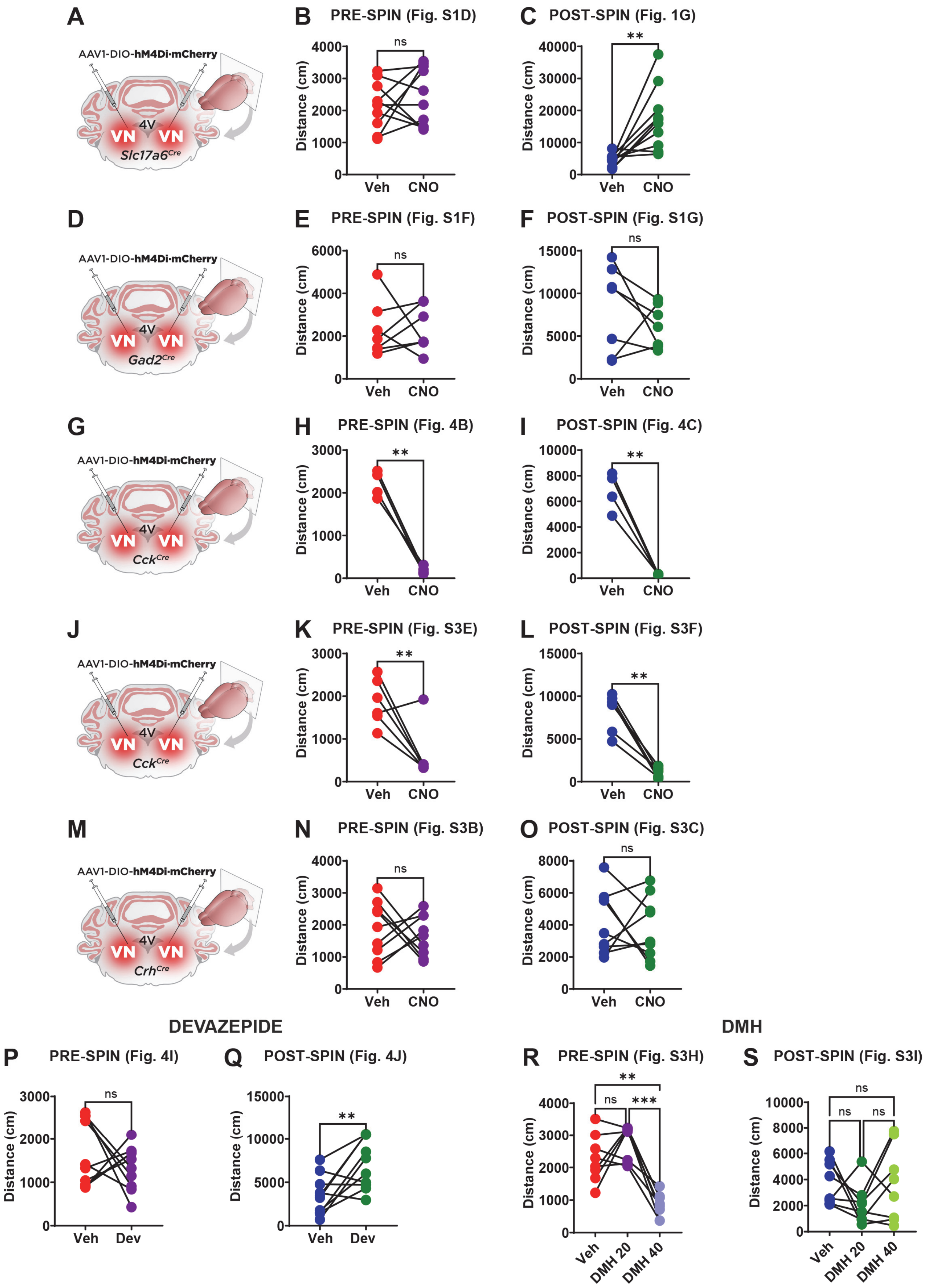

A
AAV1-DIO-hM4Di·mCherry
4V
VN VN
Slc17a6Cre
B PRE-SPIN (Fig. S1D)
ns
Distance (cm)
Veh CNO
C POST-SPIN (Fig. 1G)
**
Distance (cm)
Veh CNO
D
AAV1-DIO-hM4Di·mCherry
4V
VN VN
Gad2Cre
E PRE-SPIN (Fig. S1F)
ns
Distance (cm)
Veh CNO
F POST-SPIN (Fig. S1G)
ns
Distance (cm)
Veh CNO
G
AAV1-DIO-hM4Di·mCherry
4V
VN VN
CckCre
H PRE-SPIN (Fig. 4B)
**
Distance (cm)
Veh CNO
I POST-SPIN (Fig. 4C)
**
Distance (cm)
Veh CNO
J
AAV1-DIO-hM4Di·mCherry
4V
VN VN
CckCre
K PRE-SPIN (Fig. S3E)
**
Distance (cm)
Veh CNO
L POST-SPIN (Fig. S3F)
**
Distance (cm)
Veh CNO
M
AAV1-DIO-hM4Di·mCherry
4V
VN VN
CrhCre
N PRE-SPIN (Fig. S3B)
ns
Distance (cm)
Veh CNO
O POST-SPIN (Fig. S3C)
ns
Distance (cm)
Veh CNO
DEVAZEPIDE
P PRE-SPIN (Fig. 4I)
ns
Distance (cm)
Veh Dev
Q POST-SPIN (Fig. 4J)
**
Distance (cm)
Veh Dev
DMH
R PRE-SPIN (Fig. S3H)
**
ns ***
Distance (cm)
Veh DMH 20 DMH 40
S POST-SPIN (Fig. S3I)
ns
ns ns
Distance (cm)
Veh DMH 20 DMH 40

**Figure S6. Between-subject effects on ambulatory behavior of chemogenetic and pharmacological interventions with or without .** A) *Slc17a6*$^{Cre}$ mice were bilaterally injected in the VN with AAV1-DIO- hM4Di-mCherry (VGLUT2$^{VN}$:hM4Di mice) to inhibit VGLUT2$^{VN}$ neurons upon CNO administration. (B) Between-subject changes in locomotor activity in *Slc17a6*$^{VN}$:hM4Di mice (in a 5-min, open-field session) 30 min after CNO or vehicle administration; (n=10; paired t-test, $P>0.05$). (C) Between-subject changes in the cumulative 1-h locomotor activity in VGLUT2$^{VN}$:hM4Di mice after spin stimulation (n=10; paired t-test, $P<0.01$). (D) *Gad2*$^{Cre}$ mice were bilaterally injected in the VN with AAV1-DIO- hM4Di-mCherry (GAD2$^{VN}$:hM4Di mice) to inhibit GAD2$^{VN}$ neurons upon CNO administration. (E) Between-subject changes in locomotor activity in GAD2$^{VN}$:hM4Di mice (in a 5-min, open-field session) 30 min after CNO or vehicle administration; (n=7; paired t-test, $P>0.05$). (F) Between-subject changes in the cumulative 1-h locomotor activity in GAD2$^{VN}$:hM4Di mice after spin stimulation (n=7; paired t-test, $P>0.05$). (G) *Cck*$^{Cre}$ mice were bilaterally injected in the VN with 0.35 µl of AAV1-DIO- hM4Di-mCherry (GAD2$^{VN}$:hM4Di mice) to inhibit CCK$^{VN}$ neurons upon CNO administration. (H) Between-subject changes in locomotor activity in CCK$^{VN}$:hM4Di mice (in a 5-min, open-field session) 30 min after CNO or vehicle administration; (n=4; paired t-test, $P<0.01$). (I) Between-subject changes in the cumulative 1-h locomotor activity in CCK$^{VN}$:hM4Di mice after spin stimulation (n=4; paired t-test, $P<0.01$). (J) *Cck*$^{Cre}$ mice were bilaterally injected in the VN with 0.2 µl of AAV1-DIO- hM4Di-mCherry (GAD2$^{VN}$:hM4Di mice) to inhibit CCK$^{VN}$ neurons upon CNO administration. (K) Between-subject changes in locomotor activity in CCK$^{VN}$:hM4Di mice (in a 5-min, open-field session) 30 min after CNO or vehicle administration; (n=6; paired t-test, $P<0.01$). (L) Between-subject changes in the cumulative 1-h locomotor activity in CCK$^{VN}$:hM4Di mice after spin stimulation (n=6; paired t-test, $P<0.01$). (M) *Crh*$^{Cre}$ mice were bilaterally injected in the VN with AAV1-DIO-hM4Di-mCherry (CRH$^{VN}$:hM4Di mice) to inhibit GRH$^{VN}$ neurons upon CNO administration. (N) Between-subject changes in locomotor activity in CRH$^{VN}$:hM4Di mice (in a 5-min, open-field session) 30 min after CNO or vehicle administration; (n=9; paired t-test, $P>0.05$). (O) Between-subject changes in the cumulative 1-h locomotor activity in CRH$^{VN}$:hM4Di mice after spin stimulation (n=9; paired t-test, $P>0.05$).(P) Between-subject changes in spontaneous ambulatory activity of mice 30 min after devazepide (DEV; 1 mg/kg) or vehicle administration in a 5-min, open-field test (n=10; paired t-test, $P>0.05$). (Q) Between-subject changes in cumulative 1-hour locomotor activity of mice 30 min after devazepide (DEV; 1 mg/kg) or vehicle administration after spin stimulation (n=10; paired t-test $P>0.01$ effect of drug). (R) Between-subject changes in spontaneous ambulatory activity of mice 30 min after dimenhydrinate (DMH; 20 mg/kg or 40 mg/kg) or vehicle administration in a 5-min, open-field test (n=8; one-way ANOVA,** $P<0.01$, *** $P<0.001$ effect of drug). (S) Between-subject changes in cumulative 1-hour locomotor activity of mice 30 min after dimenhydrinate (DMH; 20 mg/kg or 40 mg/kg) or vehicle administration after spin stimulation (n=8; one-way ANOVA, $P>0.05$ effect of drug).

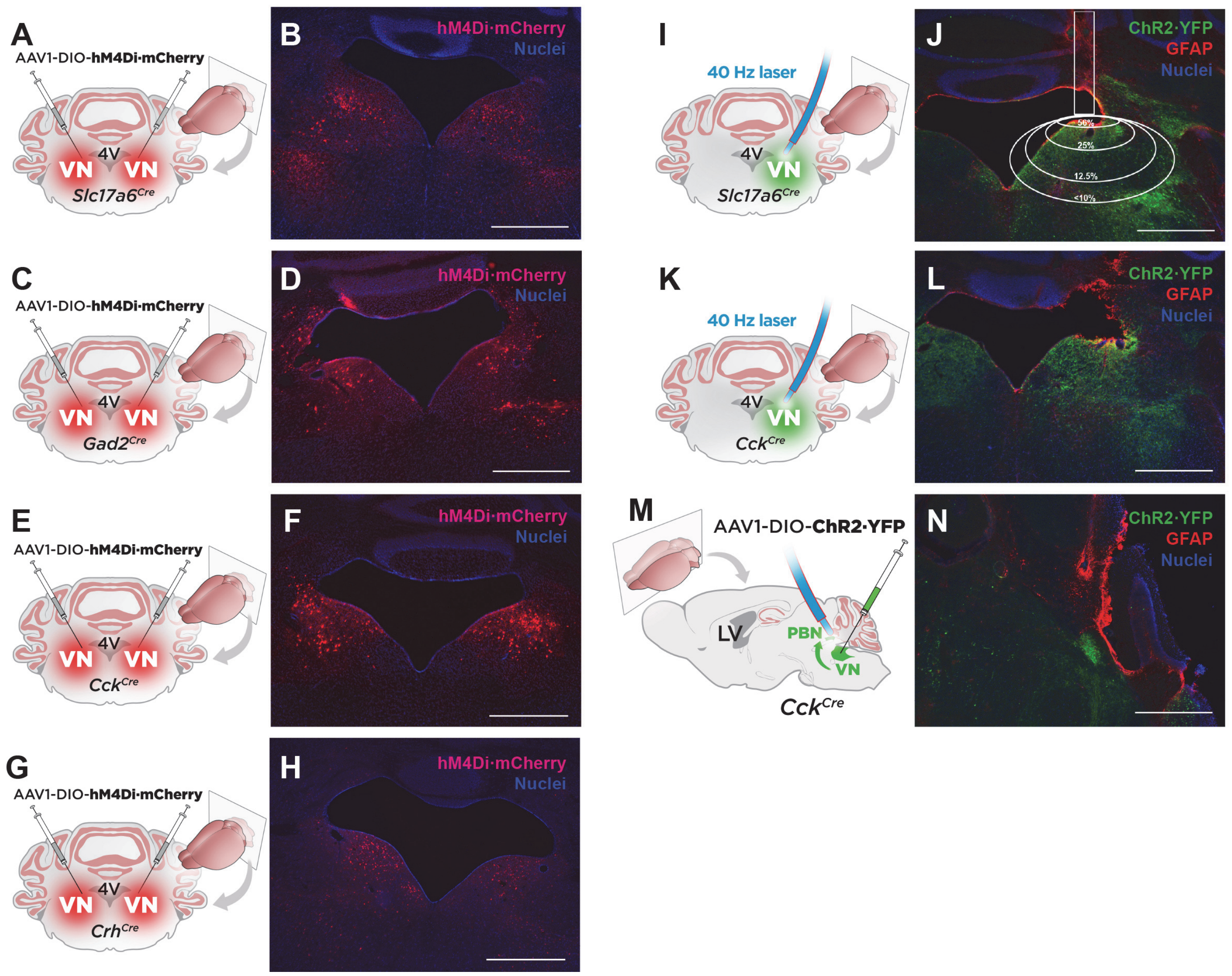


**Figure S7. Representative images of VN injection targeting and VN and PBN optic fiber implantation.** (A-B) Cartoon depicting (A) and representative targeting (B) of *Slc17a6*$^{Cre}$ mice bilaterally injected with AAV-DIO-hM4Di-mCherry in the VN. (C-D) Cartoon depicting (C) and representative targeting (D) of *Gad2*$^{Cre}$ mice bilaterally injected with AAV-DIO-hM4Di-mCherry in the VN. (E-F) Cartoon depicting (E) and representative targeting (F) of *Cck*$^{Cre}$ mice bilaterally injected with AAV-DIO-hM4Di·mCherry in the VN. (G-H) Cartoon depicting (G) and representative targeting (B) of *Crh*$^{Cre}$ mice bilaterally injected with AAV-DIO·hM4Di-mCherryin the VN. (I-J) Cartoon depicting (I) and representative targeting (J) of *Slc17a6*$^{Cre}$ mice unilaterally injected with AAV-DIO-ChR2·YFP in the VN and with fiber implantation in the same region. White rectangle depicts position of optic fiber tip. White circles indicate percentage of the light intensity compared to the power at the fiber tip. (K-L) Cartoon depicting (K) and representative targeting (L) of *Cck*$^{Cre}$ mice unilaterally injected with AAV-DIO-ChR2·YFP in the VN and with fiber implantation in the same region. (M-N) Cartoon depicting (M) and representative projection targeting (N) of *Cck*$^{Cre}$ mice unilaterally injected with AAV-DIO-ChR2·YFP in the VN and fiber implantation in the PBN. Scale bar: 800 µm

**Table S1. Mapping of monosynaptic vestibular projection targets.**

| | **AIBS Mouse Connectivity** | | **Machuca-Márquez et al.** | | |
|---|---|---|---|---|---|
| | C57Bl/6J 121146455 | *Slc17a6*-Cre 300687607 | *Slc17a6*-Cre | *Cck*-Cre | *Gad2*-Cre |
| **VN** | ++++ | ++++ | ++++ | +++ | ++ |
| **PBN** | ++++ | +++ | ++++ | +++ | ++ |
| **GRN** | +++ | ++ | ++ | +++ | ++ |
| **Cer IX/X** | +++ | ++ | +++ | + | - |
| **FN** | ++ | ++ | ++ | +/- | - |
| **NTS** | ++ | + | ++ | +/- | + |

AIBS: Allen Institute for Brain Science; VN: Vestibular nucleus; PBN: Parabrachial nucleus; GRN: Gigantocellular reticular nucleus; Cer IX/X: Cerebellar lobules IX/X; FN: Fastigial nucleus; NTS: Nucleus tractus solitarius.
